# Native-state proteomics of Parvalbumin interneurons identifies novel molecular signatures and metabolic vulnerabilities to early Alzheimer’s disease pathology

**DOI:** 10.1101/2023.05.17.541038

**Authors:** Prateek Kumar, Annie M Goettemoeller, Claudia Espinosa-Garcia, Brendan R. Tobin, Ali Tfaily, Ruth S. Nelson, Aditya Natu, Eric B. Dammer, Juliet V. Santiago, Sneha Malepati, Lihong Cheng, Hailian Xiao, Duc Duong, Nicholas T. Seyfried, Levi B. Wood, Matthew JM Rowan, Srikant Rangaraju

## Abstract

One of the earliest pathophysiological perturbations in Alzheimer’s Disease (AD) may arise from dysfunction of fast-spiking parvalbumin (PV) interneurons (PV-INs). Defining early protein-level (proteomic) alterations in PV-INs can provide key biological and translationally relevant insights. Here, we use cell-type-specific in vivo biotinylation of proteins (CIBOP) coupled with mass spectrometry to obtain native-state proteomes of PV interneurons. PV-INs exhibited proteomic signatures of high metabolic, mitochondrial, and translational activity, with over-representation of causally linked AD genetic risk factors. Analyses of bulk brain proteomes indicated strong correlations between PV-IN proteins with cognitive decline in humans, and with progressive neuropathology in humans and mouse models of Aβ pathology. Furthermore, PV-IN-specific proteomes revealed unique signatures of increased mitochondrial and metabolic proteins, but decreased synaptic and mTOR signaling proteins in response to early Aβ pathology. PV-specific changes were not apparent in whole-brain proteomes. These findings showcase the first native state PV-IN proteomes in mammalian brain, revealing a molecular basis for their unique vulnerabilities in AD.

## INTRODUCTION

A major goal in cellular neuroscience is to elucidate how the molecular signatures of unique neuronal subtypes translate to their functional diversity in intact circuits. Single-neuron transcriptomic studies have recently provided unparalleled access to the genetic diversity of dozens of unique brain cell classes^1^. Functional information is nonetheless limited in these transcriptomic studies, due to substantial discordance between mRNA and protein levels, especially in neurons^2–4^. More direct proteomic studies relying on physical isolation of individual neuron types are also inadequate, as physical isolation of individual neurons is poorly tolerated, and of those that do survive, the vast majority of their functional surface area (i.e., dendrites and axons) is lost^5, 6^. To overcome these limitations, we recently developed an *in vivo* strategy called cell type-specific in vivo biotinylation of proteins (CIBOP). When coupled with mass spectrometry, CIBOP can resolve native state proteomes from physically unaltered cell subtypes *in vivo*^7^. Key technical advancements, especially relating to neuronal subtype-specific targeting across different disease models, are also necessary to fully realize the potential of this method via extension to distinct classes of excitatory and inhibitory neurons. The recent discovery of highly versatile enhancer-AAVs^8^ have the potential to fulfill these requirements, with tools targeting inhibitory interneurons receiving major initial development^9, 10^.

Inhibitory interneurons account for 10-20% of neurons in the brain^11^. Alterations in inhibitory interneuron function appear responsible for circuit and behavioral dysfunction in several neurological diseases. In particular, dysfunction of fast spiking, parvalbumin-expressing interneurons (PV-INs) are implicated in epilepsy, neurodevelopmental, and neurodegenerative diseases including Alzheimer’s disease (AD)^12, 13^, a likely consequence of their role in maintaining circuit excitability locally, and brain state more generally, coupled with their substantial energy requirements^14^. Together, this cell class represents a promising locus for designer treatments across several major neurological disorders. Therapeutic failures are common in brain diseases, potentially due to unpredictable competing cell-type-specific responses. Thus to enhance future therapeutic efficacy, high-resolution native state proteomic signatures of individual cell classes in wild type and disease models may be required. Therefore, we implemented a versatile, systemic AAV-CIBOP intersectional approach^7, 9, 15^ to characterize and compare native state *in vivo* PV-IN proteomes from both wild type mice and in a mouse model of early AD pathology. A novel enhancer-AAV targeting method was used to express Cre recombinase specifically in PV neurons throughout the cortex and hippocampus of Rosa26^TurboID^ mice^7^. Upon Cre-mediated recombination, TurboID was expressed selectively in PV-INs, leading to robust cellular proteomic biotinylation. This PV-IN CIBOP approach identified over 600 proteins enriched in PV-INs, including canonical proteins as well as over 200 novel PV-IN proteins. The PV-IN proteome was enriched in mitochondrial, metabolic, ribosomal, synaptic, and a large number of neurodegeneration genetic risk proteins, suggesting unique vulnerabilities of PV-INs in AD.

AD is arguably the most impactful and intractable neurodegenerative disease worldwide. Interestingly, selective alterations in PV-IN physiology are increasingly appreciated across several distinct *in vivo* models of AD pathology^16–18^ which may contribute to prolonged cognitive dysfunction arising during the long-lasting, early stages of the disease^17, 19, 20^. Using network analyses of human post-mortem brain proteomes from controls and AD cases, we first identified a PV protein-enriched co-expression module (M33) strongly associated with cognitive resilience in a longitudinal aging study. Based on these lines of evidence, we next extended PV-IN CIBOP to a mouse model of hAPP/Aβ pathology^21, 22^. We found that PV-INs in pre-plaque (3 month old) 5xFAD mice exhibited extensive alterations in their mitochondrial and metabolic, cytoskeletal, and synaptic proteins, coinciding with decreased Akt/mTOR signaling. Several of these changes were validated using optogenetics, patch clamp, and cell-type-specific biochemistry. Strikingly, many of the proteomic changes noted in PV-INs in response to early hAPP/Aβ pathology were not resolved in the bulk brain proteome, suggesting that these cell-type-specific alterations are largely non-overlapping in early AD.

Overall, our studies using the CIBOP approach reveal novel native-state proteomic signatures and identify potential molecular vulnerabilities of PV-INs to neurodegeneration in AD, and nominate potentially high-value targets otherwise hidden in the bulk proteome. This enhancer AAV-CIBOP strategy will also be broadly applicable to understanding molecular complexity of PV-INs and other neuronal subtypes, across any mouse model of health or disease.

## RESULTS

### Native-state proteomics of PV interneurons in mouse brain

To achieve proteomic biotinylation of PV-INs in their native state in vivo, we employed the recent CIBOP strategy in combination with a PV-IN-specific enhancer-targeting AAV^9^, modified to drive expression of Cre recombinase (referred to here as ‘E2.Cre’).The E2.Cre construct (which also contained a 2A.GFP sequence following the Cre insert) was packaged into an AAV serotype (PHP.eB) optimized for brain-wide expression following retro-orbital (RO) sinus injection^15^. RO injections of AAV.E2.Cre were then performed in Rosa26^TurboID/wt^ (henceforth referred to as PV-CIBOP) or wild type (WT) control mice (**Fig 1A**). In a subset of studies that incorporated acute slice electrophysiology, a 2^nd^ AAV construct containing a floxed TdTomato sequence was simultaneously co-injected in the same sinus to assist with fluorescent-targeted patch clamp, because the GFP fluorescence was not visible in ex vivo slices. Starting three weeks after RO injections, mice were provided with biotin-supplemented water continuously for 2 weeks prior to euthanasia^7^.

**Figure 1.**
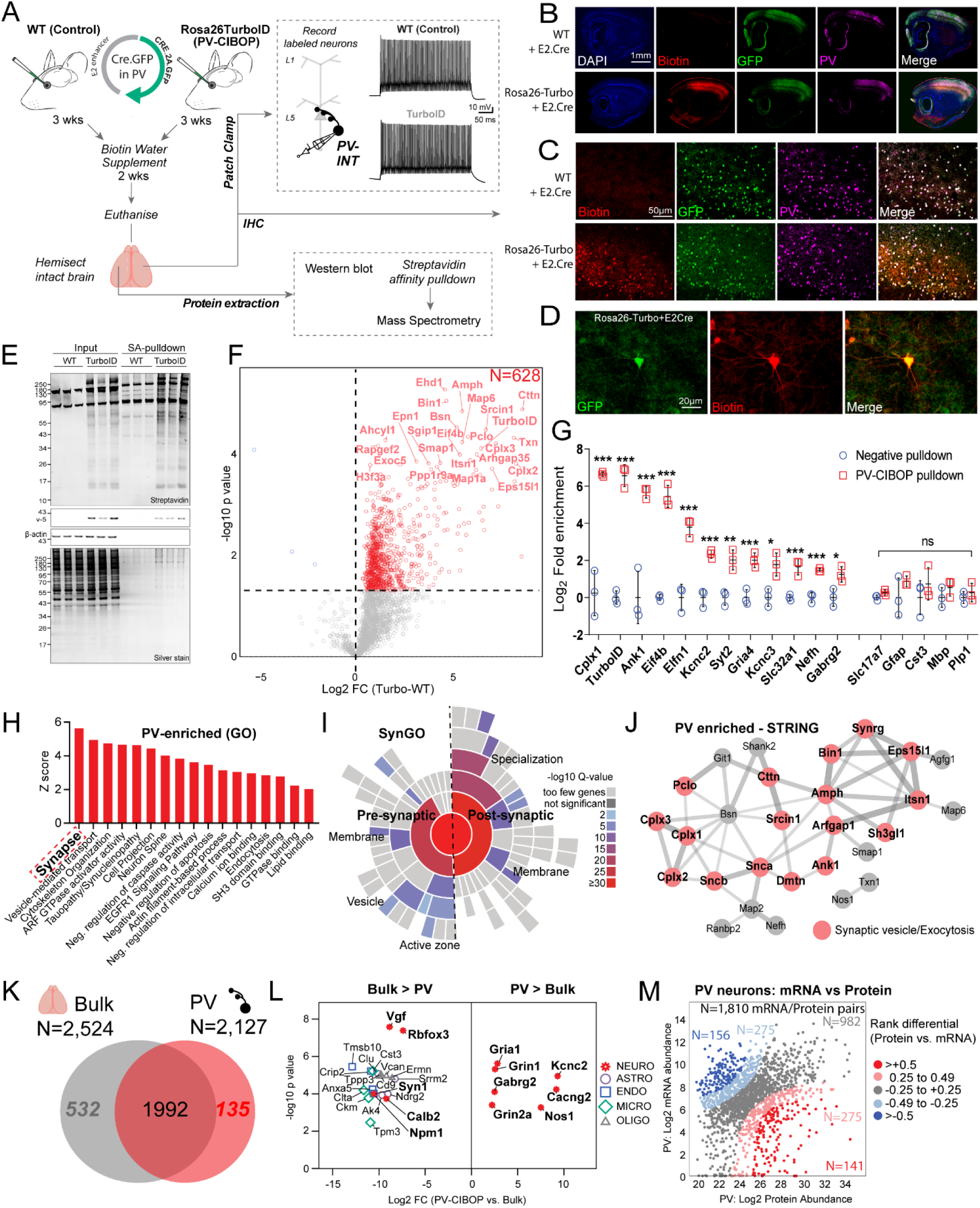
Native-state proteomics of PV-INs by CIBOP. **A.** Experimental approach: E2 enhancer Cre AAV was retro-orbitally delivered to WT (Control) or Rosa26TurboID/wt (PV-CIBOP) mice (n=3 per genotype, including males and females) followed by 3 weeks of Cre-mediated recombination, and 2 additional weeks of biotin supplementation (drinking water). The brain is then prepared for cortical slice electrophysiology, immunohistochemistry (IHC) and biochemical studies. **B-D.** IHC of fixed brain sections confirmed biotinylation (red) in PV-INs (Pvalb: green) in the cortex (Ctx) and hippocampal (HC) regions of PV-CIBOP but not control mice (B: 4x and C: 20x magnification; D: Higher magnification (60x) images from HC and Ctx are shown). **E.** Top: Western Blot (WB) of input (bulk brain tissue homogenates) and streptavidin affinity purification (pulldown) samples confirms strong protein biotinylation in PV-CIBOP (labeled) as compared to limited biotinylation (endogenously biotinylated proteins) in control animals. Bottom: Silver stained gels of inputs and pulldown samples corresponding to WB images above. **F.** Volcano plot representation of differential abundance analysis of LFQ-MS data obtained from streptavidin pulldown samples, from PV-CIBOP and control mice. Red dots represent proteins biotinylated in PV-INs as compared to control mice. Most highly labeled PV-IN proteins (including TurboID) are highlighted. **G.** Top PV-enriched proteins are shown on the left (including TurboID, Cnk1, Kcnc2, Kcnc3, Erbb4, Slc32a1 and GABA-ergic proteins). In contrast, non-neuronal (Mbp, Gfap, Aldh1l1, Cotl1) and excitatory neuronal (Slc17a7) proteins were not enriched (unpaired two-tailed T-test *p<0.05, **p<0.01, ***p<0.005). **H.** Gene Ontology (GO) analyses if PV-enriched proteins (as compared to whole brain proteome lists show enrichment of synaptic vesicle, GTPase binding, cytoskeletal and cell projection related proteins. **I.** SynGO analysis of PV-enriched proteins reveals labeling of synaptic proteins in both pre and post-synaptic compartments. **J.** STRING analysis of PV-enriched proteins (>16-fold enriched over control) shows synaptic vesicle and exocytosis related proteins including complexins, ankyrins, synucleins. **K.** Venn Diagram representing degree of overlap between proteins enriched in PV neurons, with whole brain proteomes from matched animals. While majority of PV-enriched proteins were also identified in whole brain proteomes, 135 proteins were only identified in PV neurons. **L.** Top proteins differentially enriched in PV-INs as compared to the whole brain bulk proteome and those enriched in the bulk as compared to PV-INs, are highlighted. **M.** Analysis of protein vs mRNA concordance in PV-INs, using PV-enriched proteins identified by PV-CIBOP and existing single nuclear transcriptomic data from the entire class of adult mouse PV-INs (Allen Brain Atlas). Based on differentials in rank abundances (protein vs. mRNA), discordant and concordant protein/mRNA pairs are highlighted. Also see Supplemental Figures S1, S2, S3 and Supplemental Datasheet 1 for related analyses and datasets.

Acute slice current clamp recordings from fluorescently labeled neurons confirmed selective targeting and unaltered physiology of fast-spiking PV-INs by PV-CIBOP (**Fig 1A, Supp Figure S1**). To assess potential impacts of PV-specific TurboID expression and proteomic biotinylation on PV neuron function and overall local circuit activity, we obtained voltage and current clamp recordings from layer 5 pyramidal neurons and PV interneurons, respectively, in both WT control and PV-CIBOP mice (**Supp Fig S1A,B,H**). Fast, spontaneous excitatory and inhibitory synaptic events (EPSCs and sIPSCs) were isolated during pyramidal cell recordings (**Supp Fig S1B**). The amplitude, frequency and kinetic properties of both sEPSCs and sIPSCs were unperturbed in PV-CIBOP brains (**Supp Fig S1C-G**). These results indicate that PV CIBOP does not affect basal circuit excitability, and also suggests a lack of effect on synaptic receptor distributions or properties. In the same slices, neighboring epifluorescence-identified (TdTomato+) neurons were also selected for whole-cell current clamp recordings. As expected, TdTomato+ neurons exhibited fast, non-accommodating firing with narrow action potentials (**Supp Fig S1H-J**) and passive properties characteristic of fast-spiking PV interneurons (**Supp Fig S1K-M**). Coupled with our IHC findings, the fast-spiking phenotype of TdTomato+ neurons confirmed specific targeting of PV-INs using our systemic enhancer-AAV PV-CIBOP approach.

Importantly, no differences in AP firing, various biophysical features, or passive properties were observed in PV interneurons when comparing PV-CIBOP with WT controls (**Supp Fig S1I-M**). Immunohistochemical (IHC) studies from fixed tissues showed widespread biotinylation of PV positive (PV+) neurons and GFP positive neurons across the cortex, in PV-CIBOP mice, with high efficiency of labeling (**Fig 1B-C**). In contrast, minimal biotinylation was observed in WT controls. Biotinylation of PV-INs was detected in all compartments of PV-INs, spanning somatic and axo/dendritic compartments (**Fig 1D**). Moreover, we did not observe biotinylation in non-neuronal cells and observed no morphological differences in microglia or astrocytes in PV CIBOP mice (**Supp Fig S2**). Western blots (WB) from whole brain lysate excluding cerebellum showed strong biotinylation of a wide array of proteins in PV-CIBOP mice (**Fig 1E**). Conversely, few endogenously biotinylated proteins were observed in WT control lysates^7^. In addition, the TurboID insert in Rosa26-TurboID mice contains a N-term V5 tag, thus serving as a surrogate of Cre-mediated TurboID expression. We detected V5 in PV-CIBOP mice but not WT controls, confirming the validity of the PV-CIBOP approach (**Fig 1E**).

Based on successful TurboID-mediated proteomic biotinylation in PV neurons using the PV-CIBOP approach, we next enriched biotinylated proteins from PV-CIBOP and WT control brain samples to obtain PV-specific proteomic signatures by quantitative mass spectrometry (MS). Streptavidin (SA) beads were used to enrich biotinylated proteins followed by silver stain and WB (**Fig 1E**). These confirmed enrichment of biotinylated proteins (as well as V5, a surrogate of TurboID expression) from PV-CIBOP mice, as compared to minimal endogenously biotinylated proteins from WT controls, and mirrored labeling patterns observed in bulk brain lysates (inputs). Label-free quantitative MS (LFQ-MS) was performed on bulk brain lysates and SA-enriched biotinylated samples from PV-CIBOP and WT mice and 2,534 proteins were quantified (**Fig 1F, Supp Datasheet 1**). LFQ-MS of SA-enriched samples identified 628 proteins enriched in PV-CIBOP samples as compared to WT (non-biotinylated) controls (≥2-fold increased and unadjusted p value ≤0.05, including 192 proteins ≥2-fold enriched at the FDR≤0.05 threshold) (**Fig 1F**). These included canonical PV-IN proteins, such as Kv3 channels (Kcnc2, Kcnc3), Gria4, Syt2 and Ank1, while markers of excitatory neurons (Slc17a7), astrocytes (GFAP), microglia (Cst3), oligodendrocytes (Mbp, Plp1) were not enriched (**Fig 1G**) verifying the PV-IN specificity of proteomic labeling^16, 23–25^. Few proteins were preferentially enriched in the SA-enriched samples from WT mice, while the majority showed higher levels in PV-CIBOP mice, consistent with biotinylation in PV-CIBOP animals but not in non-CIBOP controls. Gene set enrichment (GSEA) (**Fig 1H**) and SynGO (**Fig 1I**) ^26^ analyses of the PV-IN proteome showed over-representation of gene ontologies (GO) including synaptodendritic and axonal localization, neurotransmission, vesicle function, synapse organization, AFT-GTPase signaling, growth factor receptor signaling, proteins involved in tauopathies and synucleinopathies, and included both pre and post-synaptic compartments. Some of the most abundant synaptic PV-IN proteins were involved in synaptic vesicle trafficking, fusion and exocytosis and included complexins (Cplx 1-3), synucleins alpha and beta (Snca, Sncb), Bin1 and amphiphysin (**Fig 1J**)^27^.

Although the majority of proteins identified in PV-CIBOP proteomes were also detected in bulk brain proteomes from these mice, we identified several proteins that were unique to or highly-enriched in either bulk or PV-IN proteomes (**Fig 1K**). Proteins only identified in PV-IN proteomes including GABAergic channels and Kv3 channel Kcnc2 as well as proteins involved in synaptic vesicle function, endocytic/endosomal pathways, GTPase binding proteins, cytoskeletal proteins (Ank1) ^24, 28^ and ribosomal large subunit proteins (Rpl15, Rpl18, Rpl24). In contrast, non-PV neuron proteins and glial proteins were preferentially enriched in the bulk brain proteome (**Fig 1L**). We also assessed the level of concordance between protein and mRNA abundances in PV-INs, by contrasting our PV-IN proteome with published reference single cell/nuclear RNAseq PV-IN transcriptomes from mouse brain (**Fig 1M, Supp Fig S3**)^29^. Pooled data from all subtypes of PV-INs and our proteome from biotinylated PV-INs were analyzed, yielding 1,810 mRNA-protein pairs (**Supp Datasheet 1)**. The overall level of concordance was modest (Spearman’s Rho 0.27, p<0.001). Nearly 50% of mRNA/protein pairs showed reasonable level of concordance, and among these, a core set of highly abundant mRNA and proteins were identified (**Fig 1M, Supp Fig S3**), which were enriched in ontologies including oxidative-reduction processes, ATP generation, synaptic transmission, glucose metabolism, endocytosis and protein transport. Low abundance proteins with high mRNA abundance (N=156) were enriched in ontologies related to complex I respiratory chain, proteasome, cytosolic and vesicle-mediated transport terms. Proteins with high abundance but low mRNA expression included 141 proteins enriched in neurogenesis, ion channel function and transporters, cytoskeletal/cell projection and translation-related proteins.

In summary, our PV-CIBOP experiments successfully identified the proteomic signature of PV-INs that is representative of their native state in mammalian brain without any evidence of physiological disturbances. Our results also reveal proteomic phenotypes of PV-INs that are not captured by bulk brain proteomics and are discordant with mRNA-level findings.

### Proteomic signatures of PV-INs in contrast to Camk2a excitatory neurons reveal molecular signatures associated with vulnerability and cognitive resilience

Our initial analyses show that the PV-CIBOP approach can label and quantify proteins highly abundant in PV-INs without disturbing their natural physiological properties. To identify molecular characteristics and disease vulnerabilities unique to PV-INs as compared to other neuronal classes, we contrasted PV-INs with Camk2a-positive excitatory neurons, also using the CIBOP approach (**Fig 2A**)^7^. In prior work, Camk2a-CIBOP was performed in Camk2a-Cre ert2/Rosa26^TurboID/wt^ mice, and Camk2a neuron-specific biotinylated proteomes were obtained from the entire cortex and hippocampus, as performed for PV-CIBOP described here^7^. As the Camk2a and PV-IN proteomic datasets were generated using identical *in vivo* and biochemical methods, as well as similar mass spectrometry parameters, the raw MS data from both studies were searched together against the mouse Uniprot database and then analyzed to identify cell type proteomic contrasts.

**Figure 2.**
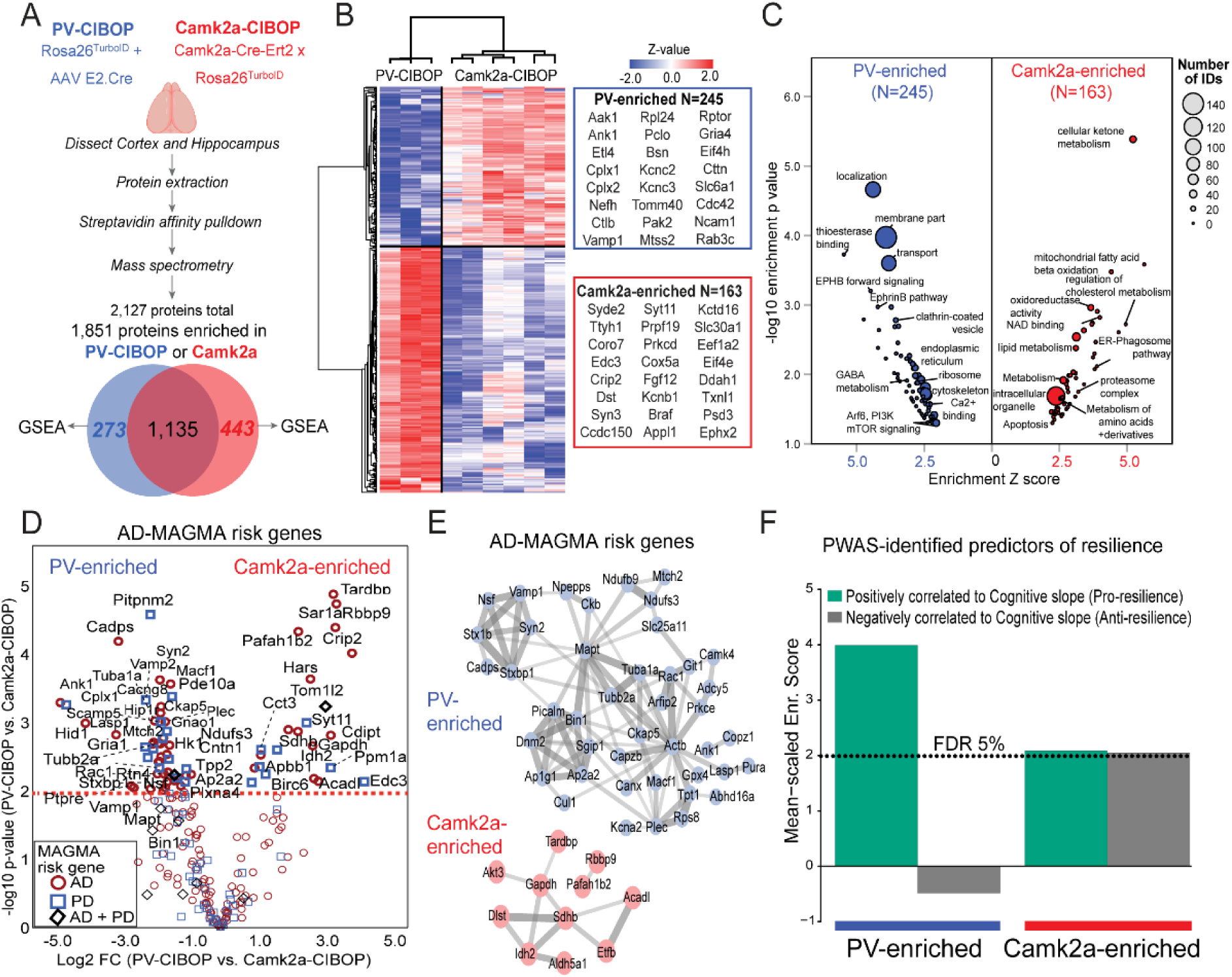
Distinct proteomic signatures and disease vulnerabilities of PV-INs and Camk2a excitatory neurons revealed by CIBOP. **A.** Experimental outline for comparative analysis of CIBOP-based proteomics of PV-INs and Camk2a excitatory neurons from the mouse cortex. Camk2a-CIBOP was achieved by tamoxifen-inducible Cre-mediated TurboID expression in Camk2a-Cre-ert2/Rosa26TurboID mice (n=6). PV-IN CIBOP was achieved by E2 enhancer AAV-mediated Cre expression (n=3), as shown in Figure 1. Non-CIBOP negative controls (n=2 per genotype) were also included. Biotinylated proteins from the cortex of Camk2a-CIBOP and PV-CIBOP as well as control mice were enriched by streptavidin pulldown, followed by label-free quantitation MS analyses. 1,841 proteins were quantified above negative samples in either PV-INs or Camk2a neurons. **B.** DEA comparing PV-CIBOP and Camk2a-CIBOP proteomes identified proteins with at least 3-fold differential enrichment (signature proteins of each neuronal class), which were then hierarchically clustered. Top proteins (based on fold-change) are shown alongside the heatmap. **C.** GSEA of PV-IN (right, blue) and Camk2a (left, red) signature proteins identified over represented terms (GO, KEGG, Wikipathways, Reactome, Pathway commons) for PV-IN and Camk2a neurons. X-axis represents enrichment Z score for a given term, and Y-axis represents level of statistical significance of enrichment. Size of each data point indicates number of protein IDs in that enrichment term. **D.** Volcano plot representation of PV-IN and Camk2a neuron signature proteins which have known genetic risk associations in Alzheimer’s disease (AD) and Parkinson’s disease (PD) based on MAGMA. Some proteins have shared genetic risk associations with AD and PD. **E.** Protein-protein interaction network (STRING) of AD-associated MAGMA risk genes that showed differential enrichment in PV-INs (Top) and in Camk2a neurons (Bottom). Clusters of mitochondrial, synaptic vesicle and endocytosis related proteins were revealed in PV-IN AD MAGMA risk genes. **F.** Enrichment of PWAS-identified proteins associated with cognitive slope in PV-enriched and Camk2a-enriched proteomic signatures. Cognitive slope was estimated in ROSMAP cases. Positive slope indicates cognitive stability or resilience while a negative slope indicates cognitive decline. Proteins positively correlated with cognitive slope are referred to as pro-resilience proteins while those negative correlated with cognitive slope are anti-resilience proteins. Enrichment of pro-resilience and anti-resilience proteins in PV-enriched and Camk2a-enriched proteins identified by CIBOP was assessed after weighting based on strength of association between proteins and cognitive slope. FDR 5% threshold is shown. Also see Supplemental Figure S4 and Supplemental Datasheet 2 for related analyses and datasets.

Of over 2,100 proteins quantified in the combined dataset, 1,841 proteins were enriched over background in either the PV-CIBOP or Camk2a-CIBOP proteomes (**Fig 2A, Supp Datasheet 2**). Of these, 1,568 and 1,408 proteins were enriched in the Camk2a-CIBOP and PV CIBOP samples, respectively, with 1,135 proteins being enriched in both. 245 proteins were highly-enriched (>4-fold) in PV-INs (including Kv3 channel proteins Kcnc2, Kcnc3) while 163 proteins were highly-enriched in Camk2a neurons (**Fig 2B**). GSEA and protein-protein interaction network analyses of these distinct neuronal proteomic signatures showed that ribosomal, GABA metabolism, ephrin B pathway, clathrin-coated vesicle, transport, cytoskeleton, endoplasmic reticulum, calcium binding, synaptic vesicle exocytosis terms as well as signaling pathways such as Akt and mTOR were over-represented in PV-IN-enriched proteins (**Fig 2C, Supp Fig S4**). In contrast, cellular metabolism, fatty acid oxidation, NAD binding, lipid metabolism, proteasome complex, ER-phagosome and mitochondrial terms were over-represented in the Camk2a-CIBOP-enriched proteome (**Fig 2C, Supp Fig S4**). Given these distinct proteomic signatures of PV-INs and Camk2a neurons, we performed upstream analyses to identify potential microRNA (miRNA) regulators of each cell type. We found an enrichment of targets of microRNAs 133a and 133b in the PV-IN proteome (**Supp Fig S4**). In an independent study that identified PV-IN and Camk2a neuron-specific miRNA signatures using miRNA tagging and affinity purification (miRAP)^30^, miRNAs 133a and 133b were highly expressed in PV-INs as compared to Camk2a neurons (**Supp Fig S4**). In our PV-IN and Camk2a CIBOP study, PV-IN proteins regulated by microRNAs 133a and 133b included synaptic vesicle and clathrin-mediated endocytic synaptic proteins (**Supp Fig S4)**. Interestingly, two microRNAs recently identified as predictors of rate of cognitive decline in humans, namely miR-29a and miR-132, were also predicted to specifically regulate PV-IN proteomic signatures^31^. These analyses suggest that molecular signatures that define PV-INs may be regulated by distinct sets of miRNAs, some of which have known associations with cognitive decline in humans.

Recent work suggests that neuron-type-specific vulnerabilities may underlie early cognitive dysfunction in neurodegenerative disorders^32^. Thus, differences in molecular signatures between distinct neuron types may underlie differential vulnerability to neurodegenerative disease. To examine this possibility, we first cross-referenced our enriched PV-IN and Camk2a CIBOP proteomic markers with known neurodegenerative disease associated risk genes from Multi-marker Analysis of GenoMic Annotation (MAGMA) analyses of human AD, Parkinson’s Disease (PD) and FrontoTemporal Dementia-Amyotrophic Lateral Sclerosis (FTD-ALS) (**Fig 2D, Supp Datasheet 2**)^33^. This integrative analysis identified 60 PV IN and 24 Camk2a-enriched proteins with known genetic causal links to AD, including genes that encode encoded proteins involved in synaptic vesicle fusion, docking and recycling (Bin1, Picalm, Dnm2, Ap1g1, Ap2a2, Sgip1), cytoskeletal and microtubule related proteins (Ank1, Actb, Tubb2a, Mapt), mitochondria proteins (Mtch2, Ndufs3, Ndufb9, Slc25a11), and SNARE complex proteins (Syn2, Stx1b, Vamp1, Nsf, Stxb1) (**Fig 2E**). In comparison, Camk2a neuron enriched AD risk genetic risk factors included proteins involved in oxidoreductase activity (Sdhb, Idh2, Aldh5a1, Etfb and Acadl) as well as the serine/threonine kinase Akt3 and TAR DNA binding protein (Tardbp) (**Fig 2E**). We also leveraged data from recent protein-wide association studies (PWAS) of post-mortem human brains from participants in the Religious Orders Study and the Rush Memory and Aging Project (ROSMAP) longitudinal study in which proteins positively correlated (n=645 pro-resilience proteins) and negatively correlated (n=575 anti resilience proteins) with cognitive slope were identified^34, 35^. As compared to the Camk2a-CIBOP proteome, we observed significantly higher enrichment of pro-resilience proteins in the PV-IN proteome (77 PV-IN-enriched vs. 36 Camk2a-enriched pro-resilience proteins), which included complexins (Cplx1, Cplx2), Ank1 and other proteins highly-abundant in PV-INs (e.g. Aak1, Cttn, Bin1, Elfn1, Bsn) in addition to ribosomal, mitochondrial, GTP binding, synaptic compartment and vesicle fusion proteins (**Fig 2F**), consistent with observations from AD-MAGMA enrichment analyses.

Together our unbiased evaluation of the proteomic signature of PV-IN, in contrast to Camk2a neurons, reveals a generalized molecular phenotype showing high translational, synaptic vesicle transport and fusion (neurotransmission), GTP binding and signaling (Akt/mTOR) activities, including many AD-related genetic risk factors and proteins associated with cognitive resilience. These indicate that the proteomic features of PV-INs may make them selectively vulnerable to pathological changes occurring in neurodegenerative diseases such as AD, and that functional or structural integrity of PV-INs may be a determinant of cognitive resilience in human brain.

### Network analyses of bulk proteomes from human post-mortem brain identify unique associations of PV-IN markers with neuropathology and cognitive dysfunction in AD

Bulk proteomics studies of post-mortem human brain from subjects with and without AD can provide important clues regarding molecular and cellular mechanisms of AD, some of which may be unique to distinct cell types in the brain, including neuronal subclasses. If PV-INs, as compared to excitatory neurons, are differentially vulnerable to neurodegenerative disease pathology in AD, network analyses of bulk brain proteomes should identify groups of co expressed proteins with distinct PV-IN versus excitatory neuronal protein marker enrichment patterns, as well as associations with clinical and pathological traits. To test this hypothesis, we interrogated MS data from published human post-mortem bulk brain proteomic studies in which co-expression network analyses identified groups (modules) of highly co-expressed proteins correlated with pathological and clinical traits of AD^36–45^. In this recent study of 516 post-mortem brain (dorsolateral pre-frontal cortex) samples from the ROSMAP and Banner Sun Health cohorts ^44, 45^, over 8,300 proteins were quantified and these coalesced into 44 co-expression modules. Samples were derived from individuals who were cognitively normal and without pathology (controls), those with asymptomatic AD (AsymAD; i.e., cognitively normal but with Aβ+tau neuropathology), and those with AD (cognitive dysfunction with Aβ+tau neuropathology)^46^. The average module-level abundances (eigenprotein) were correlated to various traits, such as amyloid burden (CERAD), neurofibrillary tangle deposition (Braak), and cognitive decline (MMSE or global cognitive function) (**Fig 3A**). GSEA previously showed that modules with increased expression in AD cases were associated with glia, metabolism, and neuroinflammation (e.g. M7) while modules with decreased expression in AD were enriched with neuronal/synaptic and mitochondrial proteins (e.g. M1, M2, M5, M11, M33), potentially indicating neuronal/synaptic loss^36, 45^. To determine whether the human brain proteomic network resolved AD mechanisms at the level of individual neuronal classes, particularly glutamatergic pyramidal neurons and GABAergic neurons (including PV, Sst, VIP subtypes), we used the Allen brain atlas of scRNAseq signatures and annotated 1,040 genes with ≥4-fold enrichment in specific neuronal classes (**Supp Datasheet 3**). Pan-glutamatergic and excitatory markers (e.g. Camk2a, Slc17a7) were enriched in modules M1, followed by M5, M22, M10 and M4. Pan inhibitory neuron proteins (e.g. Gad1 and 2, Slc32a1) were enriched in modules M33 and M23. Among the inhibitory neuron modules, M33 showed enrichment in PV-IN proteins, including PVALB and KCNC2 [Kv3.2]), while VIP interneuron markers were enriched in M23 (**Fig 3B**). Collectively, these enrichment patterns suggest that M1 (as well as M22, M5, M10 and M4) capture cellular mechanisms of AD that disproportionately impact excitatory neurons while M33 represents AD pathomechanisms impacting PV-INs.

**Figure 3.**
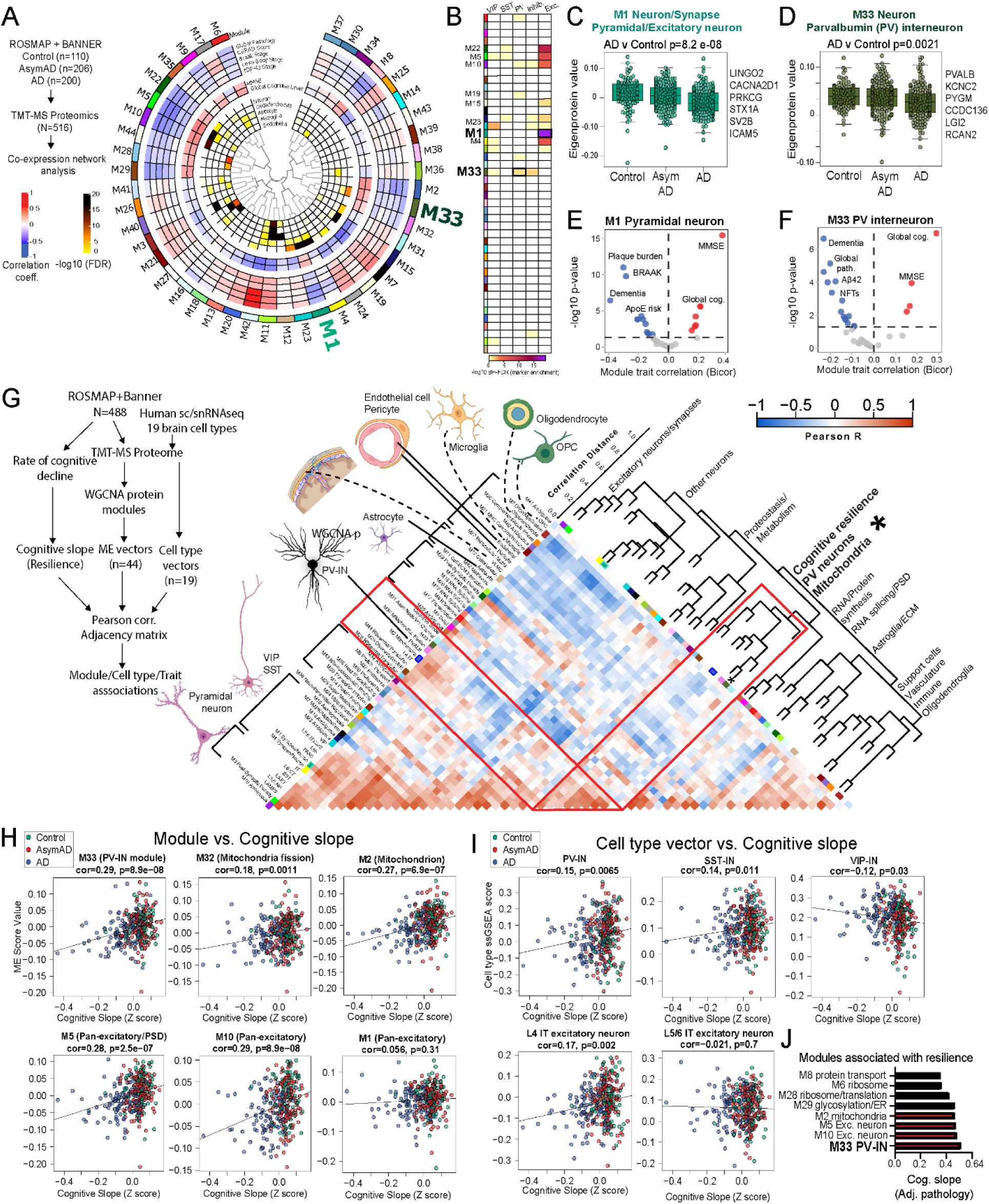
PV-IN molecular signatures are associated with neuropathological traits and cognitive resilience in analyses of bulk human post-mortem brain proteomes. **A.** Summary of network-based analysis of human bulk brain proteomes derived from post mortem frontal cortex samples from controls, AsymAD and AD cases, from ROSMAP and BANNER cohorts (adapted from Johnson et. al.)^48^. Protein co-expression modules (M1-42) are arranged in a circular manner and the central dendrogram indicates module inter-relatedness. Module trait associations are arranged in layers. Inner ring indicates associations with cell type signatures and cell type-enrichment statistical significance (-log10 FDR) are color coded. The next ring shows module-cognitive trait associations (MMSE and Global cognitive level. The third layer include module-neuropathological trait associations (red and blue indicating positive and negative correlations). Two neuron-enriched modules of particular interest to this analysis (M1 and M33) are highlighted. **B.** Modules that showed over-representation of markers of distinct classes of neurons (pan excitatory, pan-inhibitory and 3 cardinal IN classes, namely PV-IN, SST-IN and VIP-IN) are shown as a heatmap. Color indicates level of statistical significance of enrichment. M1 was enriched in pan-excitatory neuronal markers, M33 was enriched in PV-IN markers and M23 was enriched in VIP-IN markers. **C, D.** Comparisons of module abundances (eigenprotein value) of M1 (C, pan-excitatory neuronal module) and M33 (D, PV-IN module) across controls, AsymAD and AD cases. Overall ANOVA p value is shown and top neuronal class-specific proteins representative of M1 and M33 are highlighted. **E, F.** Volcano plot representations of Module-trait correlations (X-axis: Bicor, Y-axis: -log10 p-value of correlation) for M1 (E) and M33 (F). Top correlated traits (including ApoE genetic risk based on allelic combinations of ApoE ε2, 3 and 4) are labeled. See Supplemental Data x for more details. **G.** Adjacency matrix analysis based on correlations between protein co-expression modules (ME vectors), Cell type abundance vectors (based on selected markers of 19 distinct brain cell types identified by sc/snRNAseq) and cognitive slope till time of death (Left: overall outline of the analytical plan). Right: Heatmap representation of Pearson’s correlation-based adjacency matrix. Cell type vectors and module ME vectors with representative ontologies of each ME, are shown on the left of the heatmap. Dendrogram on the right indicates relatedness among ME vectors, cell type vectors and cognitive slope, and revealed a cluster of PV-IN module M33, PV IN vector, mitochondrial module M2 and M32, and cognitive slope (see Supplemental Data x for additional details). **H, I.** Associations of module eigenprotein (ME) (H) and cell type vectors (I) with cognitive slope for modules with highest positive correlations with cognitive slope. A higher cognitive slope indicates cognitive stability (or resilience) while a lower (negative) slope indicates faster cognitive decline over time. Color (reg, light blue and dark blue represent controls, AsymAD and AD cases respectively). **J.** Module eigenprotein associations with cognitive slope (resiliency) after adjustment for neuropathological features. This shows that module M33, followed by excitatory neuron modules (M5, M10) and mitochondrial modules (M2) had the highest correlation with cognitive resilience. See Supplemental Datasheet 3 for related analyses.

As individual groups, both pan-excitatory M1 and PV-IN M33 module abundances were lower in AD cases compared to control cases (**Fig 3C, D**). M1 abundance was positively correlated with cognitive function (MMSE and global cognitive function) at last clinical visit prior to death, and negatively associated with severity of dementia and neuropathology (both Aβ and tau) (**Fig 3E**). M33 abundance, like M1, was also positively associated with cognitive function and inversely associated with dementia severity and neuropathological grade (**Fig 3F**). Of note, APOE genotype (based on APOE ε4, ε3 and ε2 allelic status) was associated with excitatory (M1) but not PV-IN (M33) modules, suggesting that the genetic effect of APOE genotype on AD pathogenesis may be disproportionately linked to pyramidal neuron/excitatory rather than PV INs/inhibitory mechanisms. These protein-trait associations are limited to assessments at time of death (pathology) or last clinical evaluation prior to death, and therefore do not account for longitudinal changes in cognition occurring over several years.

To assess whether excitatory and inhibitory neuronal signatures are associated with rate of cognitive decline and the likelihood of retaining cognitive function over time, we revisited previously published protein module associations with these longitudinal cognitive trajectories from the ROSMAP study^35, 47^, and assessed the associations between cognitive slope (a measure of cognitive resilience) with abundances of protein modules as well as in silico estimates of cell type abundances (**Fig 3G, H, I**). We estimated abundances of different classes of excitatory and inhibitory neuron sub-classes along with glial and vascular cells in the brain using single-sample GSEA (ssGSEA) to determine vectors proportionate to abundances of 19 classes of human brain cell types in the human brain proteome from 598 DLPFC samples, using consensus marker lists of cell types from reference human brain single cell/nuclear RNAseq studies (**Fig 3G**)^1, 48–50^. Eigenprotein abundances of each protein co-expression module (ME vectors) were also estimated. The associations between different cell type abundance vectors with rate of cognitive decline in ROSMAP as well as with module eigenprotein vectors, was assessed, using an adjacency matrix approach based on Pearson correlation. We found that the PV-IN cell type vector, PV-IN M33 ME vector, mitochondrial M2 and M32 ME vectors as well as cognitive slope (cognitive resiliency) were significantly positively inter-correlated, identified within a PV-IN/mitochondria/cognitive resilience cluster (**Fig 3G**). In addition to PV-INs, the layer 4 IT neuronal vector was also positively correlated with cognitive slope (**Fig 3H, I**). These abundances of these two cell types were therefore the mostly likely linked to cognitive resilience. In contrast, excitatory pyramidal neurons such as layer 5/6 IT and CT neurons, were weaker correlates of cognitive trajectory. To determine whether these associations with cognitive resilience were influenced by coexisting neurodegenerative disease pathology, we evaluated prior results from Yu et. al. where module eigenprotein associations with cognitive resilience were assessed after adjusting for several neuropathologies (including amyloid, tau, synuclein pathology)^35^. After this adjustment, the PV-IN M33 module retained the highest association with cognitive resilience (**Fig 3J**).

Overall, these analyses of bulk human brain proteomes (**Fig 3**) agree with enrichment of AD genetic risk factors and protein predictors of cognitive resilience in PV-IN CIBOP proteomes (**Fig 2**) and support the idea that integrity of PV-INs and/or their proteins, may be determinants of cognitive resilience in AD. However, human post-mortem brain proteomic studies are unable to resolve the impact of aging and disease progression on neuronal protein changes in AD because sampling is performed post-mortem, leading to confounding by cause of death and stage of disease. Thus, we next endeavored to test this hypothesis using bulk brain and PV CIBOP proteomics in a mouse model of Aβ pathology.

### Neuron-specific molecular signatures resolve vulnerability of PV neurons to progressive A**β** pathology in a mouse model

We analyzed TMT-MS data from 43 WT and 43 5xFAD mice (age span 1.8-14.4 mo., 50% male, 50% female), from which over 8,500 proteins were quantified (**Fig 4A**). Here we report a sub-analysis of this dataset, restricted to neuron-enriched markers identified from reference snRNAseq studies of mouse brain (**Fig 4A-C, Supp Datasheet 4**). As expected in 5xFAD mice, age-dependent increase in Aβ pathology was associated with concomitant increase in levels of Apoe, microglial proteins (Trem2, Msn, C1qb) and astrocyte proteins (Gfap) (**Fig 4A**). Using canonical markers of different neuronal classes from mouse scRNAseq reference datasets (**Supp Datasheet 4**), we identified distinct patterns of change of neuronal proteins with aging and genotype (5xFAD vs WT), as well as biological interaction between aging and genotype (**Fig 4A**). Excitatory pyramidal neuronal proteins (Camk2a and Slc17a7) showed an age dependent decrease, although genotype had no effect on this process. In contrast, Pvalb and Sst proteins (predominantly found in PV-INs and SST GABAergic INs, respectively) increased with age in WT mice, but this trend was significantly blunted by 5xFAD genotype, particularly at 6 months of age (**Fig 4B**). Kcnc3, which encodes Kv3.3 and is also highly expressed by PV-INs, showed an age-dependent decrease in 5xFAD mice while Vip, a marker of VIP-positive INs, did not show changes related to either age or genotype. We also observed a strong positive correlation between PV-IN protein levels (Pvalb and Kcnc3) and oligodendrocyte proteins involved in myelination (Plp1 and Mbp) (**Fig 4A**) suggesting potential cell-cell interaction between PV neurons and oligodendrocytes^51^.

**Figure 4.**
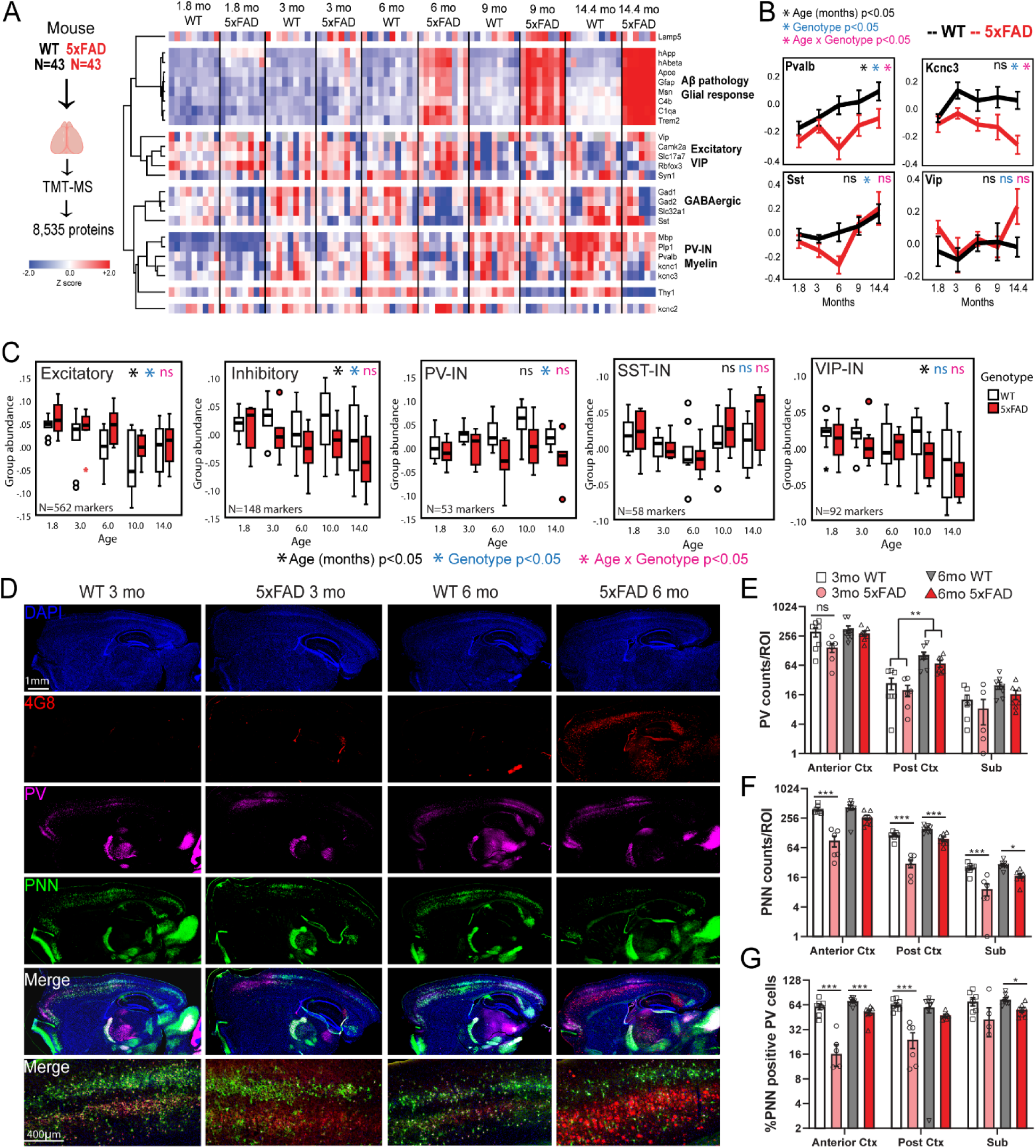
Bulk tissue proteomics of mouse brain reveals differential effects of A**β** pathology and aging on PV-INs and their peri-neuronal nets. **A.** Study outline for analysis of mouse bulk brain (cortex) TMT-MS proteomics data. 8,535 proteins were quantified by TMT-MS from 43 WT and 43 5xFAD mouse brains (including age ranges 1.8-14 months and 1:1 sex-distribution). From these, selected proteins reflective of Aβ pathology (hAβ42 peptide), resultant glial activation (Apoe, C1qa, C4b, Gfap, Trem2, Msn) and markers of excitatory and inhibitory neuronal subclasses (based on curated lists obtained from sc/snRNAseq mouse brain studies), were visualized as a heatmap after hierarchical clustering based on protein IDs. **B.** Trajectories of change in levels of PV-IN proteins (Pvalb, Kcnc3), SST-IN (Sst) and VIP-IN (Vip) based on age and genotype. Error bars represent SEM. Statistical tests included linear regression analyses including age, genotype and ‘age x genotype’ interaction terms as covariates. Levels of significance of each are indicated. **C.** Trajectories of change in overall levels of pan-excitatory, pan-inhibitory, as well as PV-IN, SST-IN, and VIP-IN proteins, based on age and genotype. We used lists of transcriptomic markers of these classes of neurons (from sc/snRNAseq datasets) that were at least 4-fold enriched in the class of interest over all other neuronal types. After normalizing and z transforming proteomic data, neuronal class-based group abundance scores were calculated and compared across ages and genotypes. Linear regression analyses were performed using age, genotype and age x genotype interaction term as covariates. Levels of significance of each, are indicated. **D.** Representative images from immunofluorescence microscopy studies of mouse brain (sagittal sections, WT and 5xFAD, ages 3 and 6 months, animals used for TMT-MS studies in A), to detect PV-INs (Pvalb protein), perineuronal nets (using WFA lectin), Aβ pathology (4G8) and DAPI. 4x tiled images and 20x images from cortex are shown. **E, F, G.** Quantitative analysis of Pvalb protein, PNNs, and proportion of Pvalb+ INs that have PNNs in the cortex (anterior and posterior to bregma) and subiculum of WT and 5xFAD mice are 3 and 6 mo (post-hoc Tukey pairwise comparisons were performed). Y axes are log2 transformed. Error bars represent SEM (Post-hoc Tukey HSD *p<0.05, **p<0.01, ***p<0.005). See Supplemental Figure S5 and Supplemental datasheet 4 for related analyses.

Using lists of markers highly expressed in excitatory or inhibitory neurons, as well as in PV-INs, SST-INs and VIP-INs from single neuronal RNAseq datasets (**Supp Datasheet 4**)^29^ which were also identified at the protein level in our proteomic dataset, we calculated composite cell-type-specific abundance scores to identify age and genotype (5xFAD vs. WT) effects in the bulk proteomic data (**Fig 4C**). When grouped, composite excitatory markers decreased with aging, and were minimally higher in abundance in 5xFAD brains as compared to WT, regardless of age. Inhibitory composite markers also showed an age-dependent decrease in abundance which became more pronounced in 5xFAD mice. Among major IN classes, PV-IN markers showed a gradual increase with age in WT mice, although this was suppressed in 5xFAD brain, consistent with the decrease in Pvalb protein (**Fig 4B**). In contrast to PV-IN marker trajectories that showed an effect of 5xFAD genotype, SST-IN and VIP-IN markers were unaffected by genotype (**Fig 4C**). Whether age-related and AD pathology-related changes in PV-IN proteins reflects protein-level changes versus alterations in cell type abundances, is unclear^52^.

To determine whether PV-IN protein changes, particularly the reduced Pvalb levels between 3-6 months of age in 5xFAD mice, are related to changes in PV-IN cell numbers, we performed IHC studies on an independent set of 3 and 6 month-old WT (n=7-8) and 5xFAD (n=7-8) brains, and assessed Pvalb protein levels along with detection of peri-neuronal nets (PNNs) by Wisteria floribunda agglutinin (WFA) in the cortex and subiculum (**Fig 4D-G**). PNNs are known to disproportionately encapsulate PV-INs in the brain and are key regulators of PV-IN excitability^53–55^. The number of Pvalb protein-positive neurons in the cortex and subiculum were not impacted by genotype (5xFAD vs WT) at 3 or 6 months of age (**Fig 4D, E, Supp Fig S5**), suggesting that changes in total Pvalb protein levels observed by MS may be related to changes at the protein level rather than changes in cell numbers. We observed decreased PNN counts in 5xFAD mice compared to WT mice, in all regions and both ages (**Fig 4F, G**). Furthermore, the proportion of PNN-positive PV-INs was also significantly lower in 5xFAD mice at both 3 and 6 months in the cortex and subiculum (**Fig 4G**).

Overall, our analyses of mouse brain proteomic data identify novel age and Aβ- dependent changes that appear to differentially impact neuronal sub-classes, where PV-INs may be selectively vulnerable in early stages of pathology in 5xFAD mice. Our histological studies suggest that observed proteomic changes in PV-IN proteins in 5xFAD brain is not explained by changes in the abundance of PV-INs^56^ although the health of PV-INs may be perturbed, as indicated by loss of PNNs around PV-INs in early stages of Aβ pathology. To better understand the molecular basis for differential vulnerability of PV-INs to AD pathology with spatiotemporal resolution, neuron class-specific native state proteomic investigations using CIBOP are warranted.

### Unique molecular signatures of PV-IN proteome in early stages of A**β** pathology *in vivo*

The native-state proteomic features of PV-INs revealed by PV-CIBOP and findings from bulk brain proteomics studies from human brain and mouse models provide indirect evidence of selective vulnerability of PV-INs to AD pathology^12^. To identify molecular events specifically occurring in PV-INs as a result of early AD pathology in their native state, we applied the AAV PV-CIBOP strategy to WT and 5xFAD mice to achieve PV-IN-specific TurboID expression, biotinylation and fluorescent labeling (**Fig 5A**). Three weeks after RO injection and 2 weeks of biotinylation, mice were euthanized at 3 months of age. This early stage of Aβ pathology (i.e., significant Aβ burden but prior to plaque formation) was chosen because this timepoint is more likely to reflect modifiable disease mechanisms of AD. IHC confirmed PV-IN-specific biotinylation in the cortex of WT and 5xFAD PV-CIBOP mice (**Fig 5B**). Flow cytometry analyses of enzymatically-dissociated PV-INs from the cortex^6^ showed that the proportion of tdTomato positivity within isolated neuronal nuclei was comparable across all groups, indicating equal efficiency of PV-IN targeting by the E2.Cre AAV approach (**Fig 5C**). Consistent with early stage of pathology, we observed minimal extracellular Aβ plaque pathology in the subiculum and cortex of 5xFAD mice, while total Aβ42 levels measured by ELISA were substantially increased in 3 month-old 5xFAD mice (**Fig 5D**). WBs of cortical lysates showed robust biotinylation and V5 protein signals in all PV-CIBOP mice (**Fig 5E**). WB of SA-enriched samples showed enrichment of biotinylated proteins in PV-CIBOP animals compared to controls, regardless of 5xFAD or WT genotype (**Fig 5F, Supp Fig S6A**).

**Figure 5.**
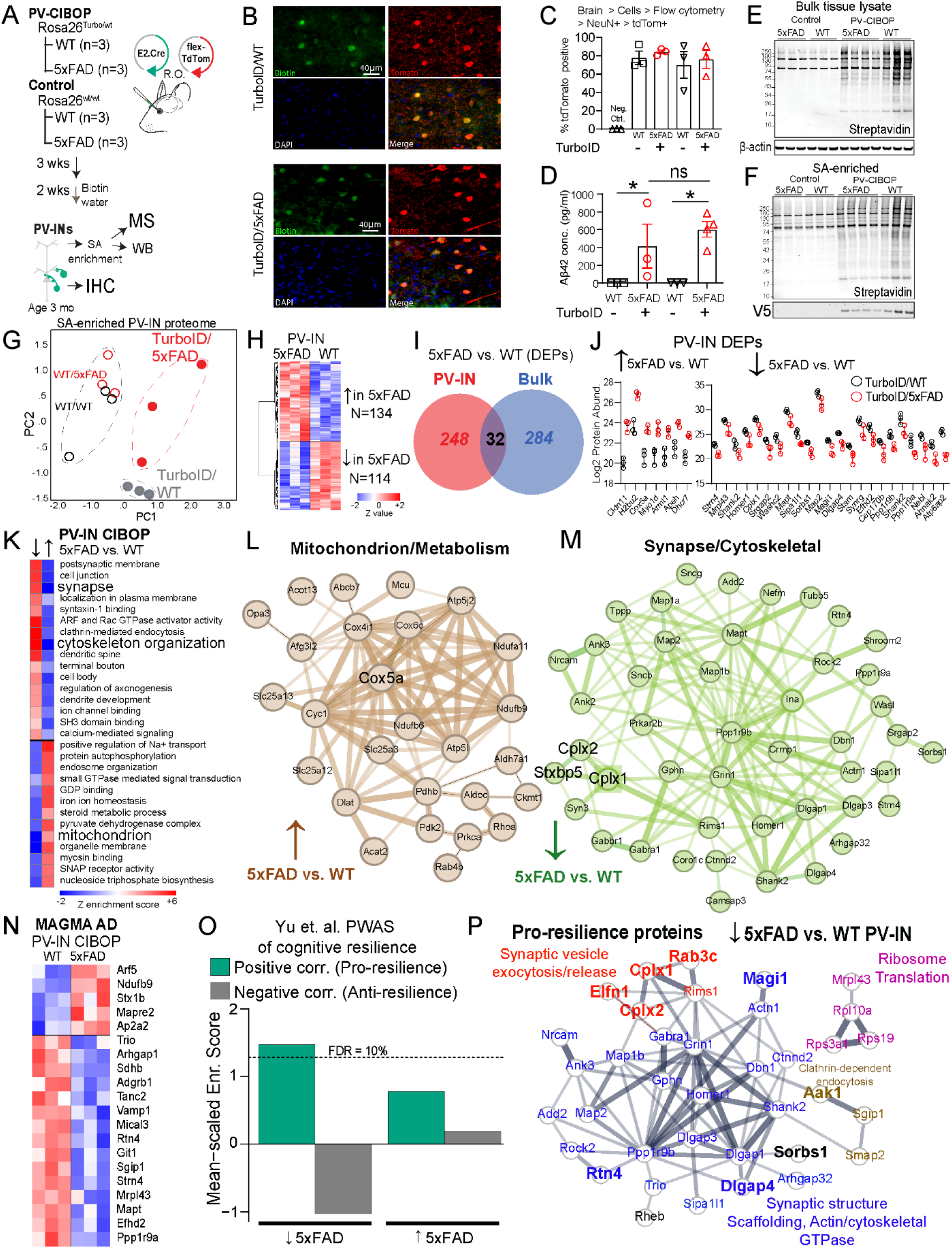
PV-IN proteomic alterations in early stages of A**β** pathology in the 5xFAD model. **A.** Experimental outline: PV-IN CIBOP was achieved via co-injection (R.O.) of E2.Cre + flex-TdTomato AAVs into TurboID mice with either WT or 5xFAD genotypes. Non-CIBOP controls involved AAV co injected non-TurboID mice (WT and 5xFAD). Mice were euthanized at age 3 months after Cre-mediated recombination and biotinylation. Tissues were used for biochemical studies (WB + MS) and IHC. **B.** IHC studies confirming PV-IN specific biotinylation in both WT and 5xFAD PV-CIBOP mice. In the cortex, majority of PV-INs (based on Pvalb immunoreactivity) were biotinylated (streptavidin) and/or expressing TdTomato. **C.** Flow cytometry analyses comparing AAV-mediated targeting efficiency of PV-INs across experimental animals. Following enzymatic digestion of cortical slices, single cell suspensions were fixed, permeabilized and labeled for live/dead indicator and NeuN. Single and live cells were further sub-gated based on NeuN positivity and proportion of tdTomato+ neurons were quantified and compared across groups. One-way ANOVA across all AAV injected groups was not significant. **D.** Aβ42 ELISA measurements (pg/mL/mg protein) from bulk brain (cortex) homogenates from all mice studied in this experiment, confirming that total Aβ42 levels were not impacted by TurboID-biotinylation status (*p<0.05, unpaired two-tailed T-test). **E,F.** WB from bulk cortical tissue lysates as well as from SA-enriched (pulldown) biotilylated fractions from all experimental mice. As compared to minimal biotinylation in control (WT and 5xFAD) mice, robust biotinylation was observed in all PV-CIBOP tissues (top). SA-enriched proteins were probed with streptavidin to confirm enrichment of biotinylated proteins in all PV-CIBOP animals. Inputs and SA enriched proteins were analyzed by LFQ-MS. **G.** PCA of bulk brain proteomes (inputs, Left) and of SA-enriched PV-IN proteomes (Right). In the inputs, minimal-to-no effects of biotinylation on bulk brain proteomes were observed. In SA-enriched proteomes (Right), all PV-IN proteomes clustered away from control samples, and further distinction was observed between 5xFAD and WT PV-IN proteomes. **H.** Heatmap representation of DEPs comparing WT/PV-CIBOP and 5xFAD/PV-CIBOP SA-enriched proteins. **I.** DEPs identified comparing 5xFAD and WT SA-enriched PV-IN proteomes minimally overlapped with DEPs identified in the bulk brain proteomes. Within the shared 32 DEPs, concordant directions of change were limited to x proteins. **J.** Top DEPs (showing at least 4-fold differential enrichment) comparing 5xFAD to WT PV-IN proteomes are shown. **K.** GSEA of DEPs comparing 5xFAD to WT PV-IN proteomes (panel G) identified clear differences based on ontology terms. While cytoskeletal, synaptic/dendritic and synaptic signaling related proteins were decreased, mitochondrial, oxphos and steroid metabolism terms were increased in 5xFAD PV-INs compared to WT PV-INs. **L, M.** STRING protein-protein-interactions (PPI) within DEPs identified in Mitochondrial (L, increased in 5xFAD PV-IN) and Synaptic/Dendritic/Cytoskeletal (M, Decreased in 5xFAD PV-IN) ontologies. Thickness of edges indicate strength of known functional or physical interactions. **N.** Heatmap representation of DEPs comparing 5xFAD to WT PV-IN proteomes, limited to proteins encoded by genes with known genetic risk associations in AD (AD-MAGMA). **O.** Analysis of enrichment of PWAS-identified proteins within DEPs (5xFAD vs. WT PV-IN proteomes) associated with cognitive resilience in PV-enriched and Camk2a-enriched proteomic signatures (as in Fig 3, pro-resilience proteins are positively correlated with cognitive slope; and anti-resilience proteins are negatively associated with cognitive slope in the ROSMAP cohort). Enrichment of pro-resilience and anti resilience proteins in PV-IN proteomes, comparing 5xFAD vs. WT PV-INs was assessed after weighting based on strength of association between proteins and cognitive slope. FDR 10% threshold is shown. **P.** STRING PPIs of PWAS-nominated proteins positively associated with cognitive resilience (pro resilience) that are decreased in 5xFAD PV-INs based on PV-CIBOP studies. Colors of proteins are based on shared functions and/or ontologies. Of these, proteins that are also selectively enriched in PV INs as compared to Camk2a neurons (from CIBOP studies in Fig 3) are highlighted (larger font, and bold). See Supplemental Figure S6 and Supplemental Datasheet 5 for related analyses.

LFQ-MS of bulk brain samples (inputs) and SA-enriched (i.e., PV-specific) samples identified 3,086 proteins and 2,149 proteins respectively (**Supp Datasheet 5**). PCA of bulk proteomes indicated minimal effects of biotinylation on the overall brain proteome (**Supp Fig 6B**). 1,973 proteins were enriched in the PV-IN proteome from both WT and 5xFAD PV-CIBOP groups as compared to negative control SA-enriched proteomes. Top PV-IN enriched proteins identified in WT/PV-CIBOP tissues in this experiment agreed with findings in the first PV-CIBOP run with moderate correlation across two independent experiments (Pearson’s R=0.61, **Supp Fig 6C**), thereby verifying reproducibility of PV-CIBOP proteomics. PCA of SA-enriched proteomes resolved differences between negative control and PV-CIBOP proteomes, and revealed further distinction between PV-CIBOP proteomes from WT and 5xFAD mice (**Fig 5G**). Thus, we next contrasted the PV-CIBOP proteomes from Turbo-expressing WT and 5xFAD mice and identified 248 differentially enriched proteins (DEPs) in PV-INs (134 increased and 114 decreased in 5xFAD, unadjusted p-value <0.05, **Fig 5H, Supp Datasheet 5**).

To establish whether PV-INs display unique cell-type-specific vulnerabilities in response to early Aβ pathology, we compared DEPs (5xFAD vs. WT) from PV-INs with those identified from the bulk proteome. Surprisingly, very little overlap was observed between DEPs in PV-INs and those found in the bulk proteome (**Fig 5I, Supp Datasheet 5**). Only 32 DEPs in the bulk tissue were identified as DEPs in the SA-enriched PV-CIBOP proteomes, and these shared DEPs also showed poor concordance (**Supp Fig S6D**). Gene set variation analyses (GSVA) identified 210 ontologies differentially enriched in the PV-IV proteome in contrast with only 16 ontologies in the bulk proteome (**Supp Datasheet 5**). Together, this indicates a lack of concordance between the effects of 5xFAD genotype on the bulk brain proteome and on PV-IN proteome, and suggests that proteomic effects occurring in specific cell classes such as PV-INs are masked in bulk tissue.

Among DEPs identified in the PV-IN proteome, top proteins showing at least 4-fold increase in 5xFAD PV-INs included Cox5a, Dhcr7 and Apeh (**Fig 5J**). Cox5a is a Complex IV mitochondrial protein involved in ATP synthesis^57^. Dhcr7 encodes 7-dehydrocholesterol reductase that catalyzes final rate limiting steps of cholesterol biosynthesis^58^. Apeh encodes acylaminoacyl-peptide hydrolase that hydrolyses terminal acetylated residues in small acetylated peptides, including degradation of monomeric and oligomeric Aβ^59^. Synaptic structural proteins including Shank2, Homer1, Map2 were conversely decreased by at least 4- fold in 5xFAD PV-INs (**Fig 5J**). GSEA as well as GSVA of DEPs in PV-IN proteomes showed that proteins involved in mitochondrial function, iron homeostasis, steroid biosynthesis, small GTPase signaling, and GDP binding were generally increased in 5xFAD PV neurons (**Fig 5K, L, Supplemental Datasheet 5**). In contrast, proteins associated with structural/cytoskeletal, synaptic, axonal and dendritic ontologies were generally decreased in 5xFAD PV-INs (**Fig 5K**). GSVA of bulk and PV-IN proteomes also verified 36 post-synaptic proteins (post-synapse GO:0098794) showed decreased levels in 5xFAD PV-INs, including structural constituents of the post-synapse (Dlgap1, Homer1, Gphn, Ina, Shank2, Git), enzymes with kinase activity or binding (Ppp1r9b, Bcr, Rtn4, Rheb, Prkar2b, Map2), neurotransmitter receptors (Grin1, Gabra1, Gabbr1), dendritic spine proteins (Grin1, Homer1, Dlgap3, Shank2, Dbn1, Bai1, Tanc2, Ncam1) and ribosomal subunits (Rps19, Rpl10a). Pre-synaptic proteins involved in synaptic vesicle fusion and exocytosis, including complexins (Cplx1, 2 and 3), showed decreased levels in 5xFAD PV-INs (**Fig 5M**). Interestingly, Cplx1 and 2 (but not Cplx3) were more also abundant in PV-INs as compared to Camk2a neurons in our CIBOP studies (**Supp Fig S6**). Several MAGMA-identified AD genetic risk factors showed differential abundances in 5xFAD PV-INs as compared to WT PV-INs (e.g., decreased Ppp1r9a, Mapt, Git1; increased Arf5, Ndufb9, Stx1b (**Fig 5N**).

To predict whether changes in the PV-IN synaptic protein landscape would result in detrimental or protective consequences, we cross-referenced the 5xFAD vs. WT PV-IV DEP list against pro-resilience and anti-resilience proteins (**Supp Datasheet 5**). We found that pro resilience proteins were over-represented while anti-resilience factors were under-represented in proteins that decreased in 5xFAD PV-INs (**Fig 5O**). The pro-resilience proteins which were decreased specifically in PV-INs in early 5xFAD pathology included synaptic structural, synaptic scaffolding, actin/cytoskeleton (Dlgap13/4, Shank2, Homer1, Dbn1, Map1b, Map2, Ank3), ribosome (Rpl10a, Rps19, Mrpl43), mTOR-C1 regulating protein (Rheb), clathrin-dependent endocytic (Aak1, Sgip1, Smap2) and synaptic vesicle fusion/exocytosis/release related proteins (Cplx1, Cplx2, Elfn1, Rab3c, Rims1) (**Fig 5P**). Many of these pro-resilience proteins that were decreased in 5xFAD PV-INs (e.g. Cplx1, Cplx2, Elfn1, Rab3c, Rtn4, Dlgap4, Sorbs1, Magi1) were also highly enriched in PV-INs as compared to Camk2a neurons (**Fig 5P, Supp Fig S6E-F**), indicating that these changes in early AD pathology may indeed be specific to PV-INs.

Overall, these mouse-human integrative analyses indicate that altered levels of PV-IN bouton and dendritic proteins at early stages of Aβ pathology may have detrimental synaptic effects, and thus represent a proteomic signature of decreased cognitive resilience in AD pathology. However, these changes may also represent an early homeostatic response more associated with resilience. Thus, we next aimed to understand the functional impact of these PV-specific synaptic proteomic alterations.

### Early A**β** pathology impacts PV-pyramidal cell neurotransmission and network activity

To evaluate the effect of early Aβ pathology on PV-to pyramidal cell neurotransmission, we again leveraged the ‘E2’ enhancer to express optogenetic actuators specifically in PV-INs (**Fig 6, Supp Fig S7**. This optogenetic approach allows for the study of PV-specific neurotransmission properties, with postsynaptic pyramidal cell recordings representing an integrated response to PV-specific, action potential-evoked neurotransmission. The red shifted opsin C1V1 (AAV.E2.C1V1) was stereotactically injected in layer 5 of S1 in both WT and 5xFAD mice. For 5xFAD experiments, AAV.Camk2.YFP was simultaneously co-injected and served to confirm accurate viral targeting (**Fig 6A**) with injections performed 1-2 weeks before procurement of acute slices in two separate age cohorts (∼2 or ∼3 months old, 7 or 14 weeks respectively) and voltage clamp recordings were obtained from postsynaptic pyramidal neurons in layer 5 (**Fig 6B**). Short (∼0.2 ms) LED-light pulses (590 nm) reliably evoked IPSCs every trial (0.1 Hz inter-trial interval) in both WT and 5xFAD recordings with minimal temporal jitter (1.5 mM extracellular Ca^2+^). The amplitude of the first IPSC amplitude was unchanged in both 2- and 3-month-old 5xFAD mice (e.g., 2-month-old WT: 133.5 ± 25.90 pA, n=10; 3-month-old 5xFAD: 163.7 ± 30.55 pA, n=7; p = 0.46, unpaired t-test), suggesting that GABA_A_ receptor availability was similar in the pyramidal cell postsynaptic membrane. As several synaptic vesicle fusion/exocytosis/release related proteins (Cplx1, Cplx2, Stx1b, Elfn1, Rab3c, Rims1) were altered in our 5xFAD PV-IN proteome, we more closely evaluated presynaptic properties (i.e., release probability) of PV-pyramidal synapses.

**Figure 6.**
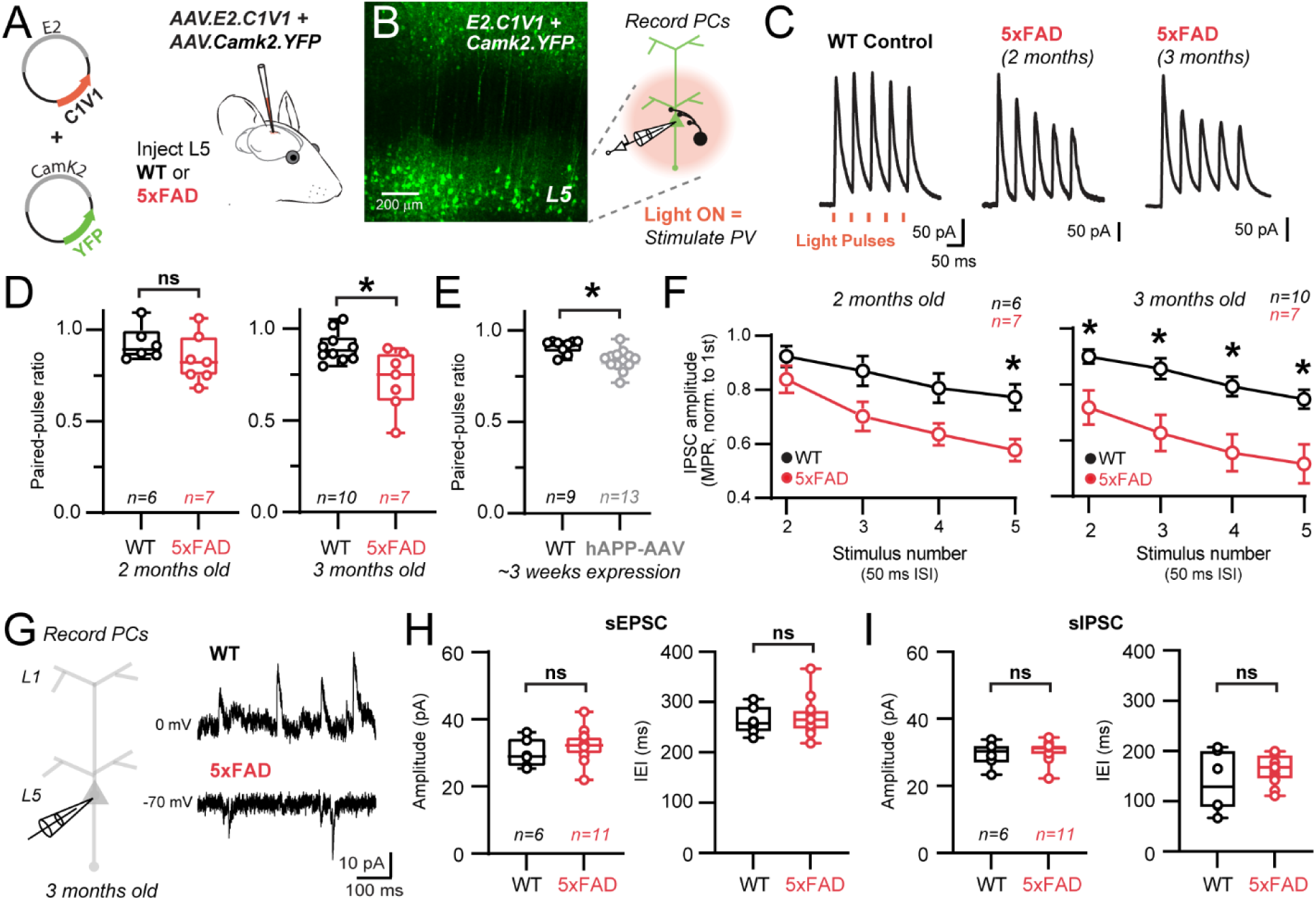
Progressive dysfunction of PV-pyramidal cell neurotransmission in young 5xFAD mice. **A.** PV-INs were targeted for optogenetic activation in WT and 5xFAD mice using E2-driven AAV expression of C1V1 in S1 cortex. AAV.Camk2.YFP was also co-injected simultaneously as a volume label in these experiments. **B.** 2-photon z-stack image showing successful targeting of L5 following ∼1 week after stereotactic surgery. Pyramidal neuron somas and apical dendrites are evident. Patch clamp cartoon depicts the experimental workflow to stimulate PV interneurons and record their synaptic properties in post-synaptic pyramidal cells. **C.** Examples of averaged voltage clamp traces from postsynaptic WT and 5xFAD pyramidal cells in layer 5 in response to short amber light pulses. PV-IN IPSCs are shown as time-locked to amber light pulses in a 20 Hz train. **D.** Quantification of paired-pulse ratios of C1V1-evoked IPSCs in pyramidal cell recordings for 2- and 3- month-old WT and 5xFAD mice (ns signifies p>0.05, unpaired two-tailed T-test) (*p<0.05, unpaired two tailed T-test). **E.** Quantification of paired-pulse ratios of ChETA-evoked IPSCs in pyramidal cell recordings ∼3 weeks following hAPP-AAV injections in L5 cortex (*p<0.05, unpaired two-tailed T-test). **F.** Quantification of changes in PV inhibitory synapse dynamics as measured via multiple pulse ratio (MPR) in both 2- and 3-month-old WT and 5xFAD mice. (*p<0.05 Two-way ANOVA with Sidak’s posthoc comparisons for each stimulus in WT and 5xFAD experiments). **G.** Voltage clamp experiments were performed examining spontaneous synaptic activity in separate cohorts of 3-month-old WT and 5xFAD mice. The holding voltage was interleaved between -70 and 0 mV throughout recordings to resolve spontaneous EPSCs and IPSCs, respectively. **H.** Quantification of spontaneous EPSC amplitude and frequency (ns signifies p>0.05, unpaired two-tailed T-test) in 3-month-old 5xFAD and WT mice. Each data point indicates an average value from all spontaneous events from individual recordings. **I.** Quantification of spontaneous IPSC amplitude and frequency (ns signifies p>0.05, unpaired two-tailed T-test) in the same recordings as in (H). Data points indicate an average value from all spontaneous events from individual recordings. See Supplemental Figure S7 for related analyses.

To examine whether modification of vesicle fusion and association proteins affected release probability and presynaptic dynamics at PV-pyramidal synapses, we measured the paired pulse ratio (PPR)^60^ and multiple pulse ratio (MPR) of optogenetically-evoked IPSCs at 20 Hz (**Fig 6C**)^61, 62^. Evaluation of the paired-pulse ratio showed modest depression (PPR ∼0.9) in WT mice at both age timepoints. In 2 month old 5xFAD mice, no difference in PPR was observed (**Fig. 6D**) although synaptic depression did intensify in 5xFAD mice during repetitive stimuli (MPR) at this age (**Fig. 6F**). In 3-month-old 5xFAD mice, stronger synaptic depression was apparent via PPR (**Fig. 6D**), and this more robust depression was now maintained throughout the stimulus train (**Fig. 6F**). Together these results indicate progressive presynaptic dysfunction in PV-pyramidal synapses following early Aβ pathology, likely involving proteins which regulate vesicular release probability. However, secondary mechanisms, such as proteins involved with vesicular docking and replenishment, axonal action potential signaling, and Ca^2+^ dynamics may also contribute. For example, changes in presynaptic parvalbumin expression may affect short-term plasticity and release probability^63^. However, we did not observe a change in bulk synaptic parvalbumin levels by MS (**Fig 4B**) or IHC **(Supp Fig S5)** at 3 months of age

We next asked whether the signature of synaptic dysfunction observed in 5xFAD mice could be recapitulated in an independent, adult-onset AD model of APP/Aβ pathology. To achieve this, we packaged the human APP gene (variant NM_000484.4) into an AAV (AAV.Ef1a.hAPP) for pan-neuronal expression (henceforth referred to as hAPP-AAV). This particular APP isoform was chosen as it has been shown to proportionally increase with aging (Koo et al. 1990, Matsui et al. 2007) and is associated with increased AD risk (Kivipelto et al. 2002, Honig et al. 2005). 5-11 week old PV-Cre mice were co-injected in S1 with hAPP-AAV and AAV.DIO.CAG.ChETA for PV-specific optogenetic control **Supp Fig S7A**. A second PV-Cre control group only received AAV.DIO.CAG.ChETA with saline. After 2-3 weeks expression, voltage clamp recordings were obtained from postsynaptic pyramidal neurons in S1 layer 5. Brief (∼4 ms) LED-light pulses (470 nm) could reliably evoke IPSCs on every trial (0.1 Hz inter trial interval) in both control and hAPP-AAV groups, with minimal temporal jitter **Supp Fig S7B,C**. Like findings with E2.C1V1 from 3-month-old 5xFAD mice, the amplitude of the first ChETA-evoked IPSC in hAPP-AAV experiments was unchanged with respect to control (Control: 68.18± 38.37 pA, n=9; hAPP-AAV: 72.70 ± 44.49 pA, n=13; p = 0.82, unpaired t-test). Also, analogous to E2.C1V1 experiments in WT mice, optogenetic stimulation with ChETA showed modest PV-pyramidal synaptic depression in controls (**Fig. 6D vs 6E**). However, similar to 5xFAD, synaptic depression measured via PPR and MPR was enhanced following hAPP AAV expression (**Fig. 6E; Supp Fig S7D**). Together these results complement our findings in 5xFAD mice, highlighting the emergence of presynaptic dysfunction at PV-pyramidal synapses following early Aβ pathology in two distinct models.

Alterations in release probability and other presynaptic dysfunction in PV-INs are expected to affect basal network excitability by disrupting excitatory/inhibitory balance. Thus, we next examined whether changes in the amplitude and frequency of spontaneous EPSCs and IPSCs were apparent in pyramidal cell recordings from 3 month old 5xFAD mice (**Fig. 6G**). Interestingly, no changes were observed in either the amplitude or frequency of excitatory and inhibitory spontaneous synaptic events (**Fig. 6 H, I**). This overall lack of network effect echoes recent work from 3-month-old 5xFAD mice in the hippocampus, where local circuit behavior and oscillations were also largely resilient to change^64^. The extensive cell-type-specific proteomic alterations we observed in PV-CIBOP and bulk proteomes may thus be reflective of early homeostatic responses seeking to maintain PV-IN and overall circuit functionality. This process would necessarily induce some degree of metabolic stress^65^. Therefore, we next sought to better understand the extent of changes to mitochondrial proteins and associated metabolic pathways specifically in PV interneurons following early Aβ pathology.

### Evidence for extensive mitochondrial protein changes in PV-INs in response to early A**β** pathology

We focused our analysis on the 300 mitochondrial proteins (annotated by MitoCarta 3.0, mouse list of 1,140 proteins) that were biotinylated in PV-INs^66^. Strikingly we identified 34 DEPs, including 30 proteins that were increased (e.g. Cox5a, Mpst, Ndufa11, Ckmt1) and 4 proteins that were decreased (Mrpl43, Septin4, Sdhb, Bphl) in 5xFAD compared to WT PV-IN proteomes (**Fig 7A**). With regards to mitochondrial pathways differentially impacted by Aβ pathology specifically in PV neurons, proteins involved in complex III, complex IV, complex V, amino acid metabolism and protein homeostasis were differentially enriched in 5xFAD PV proteomes, while mitochondrial structural (central dogma), complex II and detoxification related proteins were unaffected (**Fig 7B**). Overall, these findings are consistent with increased mitochondrial mass specifically in PV-INs in 5xFAD as compared to WT mice, rather than specific mitochondrial complex or compartment-specific effects. In contrast to the PV-IN proteome, only 26 mitochondrial proteins were differentially enriched in the bulk brain proteome, which included only 4 shared DEPs (**Fig 7C)**. Furthermore, the overall level of concordance between bulk brain and PV-IN mitochondrial protein levels was negligible (R^2^=0.0001). The increase of Cox5a levels in 5xFAD PV-INs but not in the bulk brain tissue was also validated by Western Blot, using both bulk brain homogenates and PV-specific enriched proteins (**Fig 7D**). Cox5a, a key member of complex IV, was also quantified by MS in the bulk brain proteome of WT and 5xFAD brains from an independent study of mouse brain, also showing age-dependent decrease in 5xFAD mice (**Fig 7E**), a pattern which is in stark contrast to increased levels in PV-INs at 3 months.

**Figure 7.**
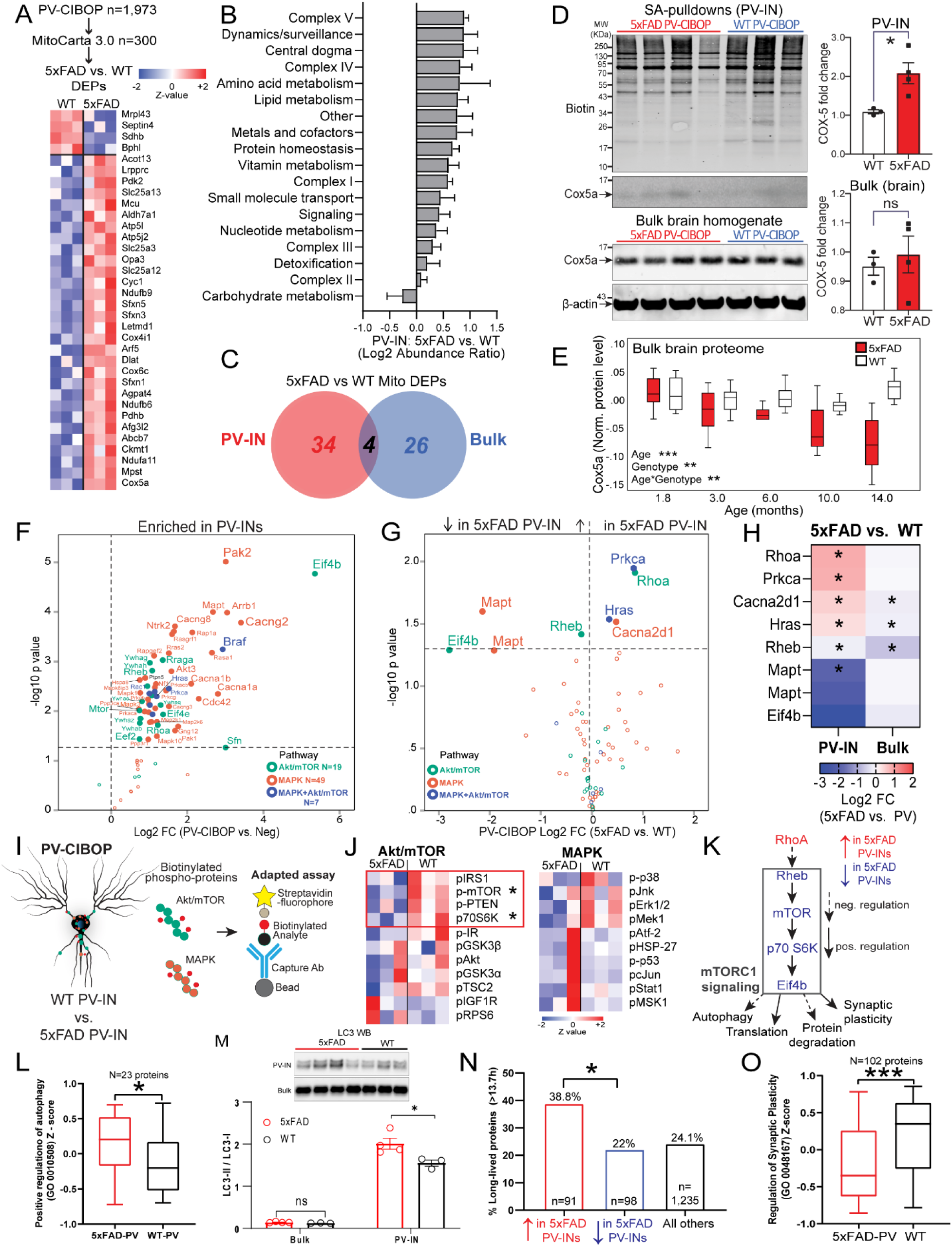
Distinct mitochondrial alterations in PV-INs at early stages of A**β** pathology. **A.** Heatmap representation of mitochondria-localized proteins that were also identified as DEPs comparing 5xFAD to WT PV-IN proteomes. Of 1,973 biotinylated proteins identified in PV-INs, 300 mitochondrial proteins were identified (based on mouse MitoCarta 3.0 list of over 1,000 mitochondrial proteins). Majority of these mitochondrial DEPs showed increased levels in 5xFAD PV-INs. **B.** Differential abundance analysis of distinct mitochondrial functional groups, comparing 5xFAD to WT PV-IN proteomes. The 300 mitochondrial proteins identified in PV-INs were categorized based on known functional and localization-related annotations (from MitoCarta 3.0). Protein levels were normalized and them group-wise abundances were estimated and compared across WT and 5xFAD genotypes (*p<0.05, **p<0.01 for unpaired two-tailed T-test, error bars represent SEM). **C.** Venn diagram of mitochondrial proteins that were identified as DEPs in either PV-IN proteomes or bulk brain cortical proteomes, comparing 5xFAD and WT mice. Minimal overlap in DEPs were observed, highlighting unique mitochondrial effects of Aβ pathology in PV-INs, not visible at the bulk tissue level. **D.** WB verification of increased Cox5a protein levels in PV-INs in 5xFAD as compared to WT mice. SA enriched pulldowns were independently performed from samples used for LFQ-MS studies. Cox5a protein band intensity was normalized to total biotinylation signal in the SA-enriched pulldowns, and to beta-actin in the bulk brain homogenates, and then compared across genotype (5xFAD vs. WT) (*p<0.05, unpaired two-tailed T-test). **E.** Cox5a protein levels, quantified by TMT-MS, from an independent set of cortical brain homogenates obtained from WT and 5xFAD mice, spanning ages 1.8 to 14 months (equal numbers of male and female mice per group, total N=86). Using linear regression modeling, age, genotype and age x genotype interaction terms were tested for associations with Cox5a protein levels. As compared to WT brain where Cox5a levels were relatively constant with aging, Cox5a levels in 5xFAD brain showed age-dependent decrease after 6 months of age. This pattern was discordant with increased Cox5a in PV-INs in 5xFAD mice at 3 months. **F.** Volcano plot representing proteins that belong to the Akt/mTOR and/or MAPK signaling pathways (curated from HGNC) that were also identified as biotinylated proteins in PV-IN-specific CIBOP proteomes. **G.** Among biotinylated Akt/mTOR and MAPK proteins in PV-INs, few were identified as DEPs comparing 5xFAD to WT PV-IN proteomes. **H.** Heatmap representation of Akt/mTOR and MAPK DEPs in PV-IN proteomes and their corresponding changes in bulk brain proteomes. PV-IN proteomic changes in these pathways were not observed at the bulk tissue level (with the exception of Rheb). **I.** Cartoon representation of adapted Luminex immunoassay to measure levels of phospho-proteins belonging to Akt/mTOR and MAPK signaling pathways. As published previously^7^, analytes of interest at first captured onto beads using a capture antibody. If this analyte is biotinylated (by PV-CIBOP), it can be detected using a streptavidin fluorophore. If not biotinylated, no signal is detected. This provides a direct estimation of phosphoproteins that reflect signaling pathway activity, with PV-IN-specific resolution. **J.** Heatmap visualization of phospho-proteins from Akt/mTOR and MAPK signaling pathways measured using the adapted Luminex assay in cortical brain lysates from WT/PV-CIBOP and 5xFAD/PV-CIBOP mice (n=3 mice per group). Adapted signal obtained from non-CIBOP brain lysates was used a background signal and subtracted from signal obtained from CIBOP adapted Luminex values. These were further normalized to TurboID abundance (as estimated by LFQ-MS from the same samples) (p<0.05, unpaired two-tailed T-test). **K.** Summary of results indicative of an overall decreased activity in mTOR signaling (particularly via mTORC1) in 5xFAD PV-INs as compared to WT PV-INs. This summary was constructed using total protein levels estimated by LFQ-MS of PV-IN proteomes, as well as phospho-protein levels by the adapted Luminex approach. mTORC1 signaling is expected to suppress autophagy, increase ribosome biogenesis and translational efficiency, decreased protein degradation and increase synaptic activity/plasticity. **L.** Comparison of proteins that positively regulate autophagy (GO 0010508) as a group, in 5xFAD and WT PV-IN proteomes (levels of 23 proteins were normalized, z-transformed and then averaged across biological replicates before group comparisons using T-test (*p<0.05). **M.** Western blot analysis of PV-IN (SA-enriched) samples and bulk brain lysates from 5xFAD and WT PV CIBOP samples, to measure LC3-II and LC3-I bands, to estimate autophagic activity. LC3-II and I bands were separately quantified and LC3-II/I ratios were calculated and compared across the two groups. Quantitative analysis is shown below (error bar represents SEM, *p<0.05, independent two-tailed T-test). **N.** Analysis of DEPs (5xFAD vs. WT PV-IN proteomes) based on published protein half-lives in mouse brain. Proteins with increased levels in 5xFAD PV-INs were skewed towards proteins with longer half lives (>13.7 days which represents the 75^th^ percentile of protein half-lives in brain). This pattern is consistent with decreased translational efficiency and/or increased protein degradation, which would disproportionately impact the relative abundances of short-lived proteins. **O.** Comparison of proteins that regulate synaptic plasticity (GO 0048167) as a group, in 5xFAD and WT PV-IN proteomes (levels of 102 proteins were normalized, z-transformed and then averaged across biological replicates before group comparisons using T-test (***p<0.005). See Supplemental Datasheet 6 for related analyses.

One interpretation of increased mitochondrial protein abundance in 5xFAD PV-INs is increased availability of mitochondrial proteins for biotinylation by TurboID, due to mitochondrial disruption during apoptosis. By design (TurboID-NES), the CIBOP approach does not access the mitochondria directly. In our studies in WT PV-CIBOP proteomes, mitochondrial DNA encoded gene products were not labeled by TurboID, while several nuclear-encoded mitochondrial proteins were biotinylated, consistent with the inability of TurboID-NES to directly access the mitochondrial compartment. Mitochondrial disruption and release of mitochondrial proteins into the cytosol, such as cytochrome C (Cytc), is a well-known marker of induction of apoptosis^67^. However, Cytc was not labeled by TurboID in both WT or 5xFAD PV-IN proteomes and was quantified at the level of noise. Given the overall pattern of increased abundances of mitochondrial proteins belonging to most mitochondrial compartments, it is likely that these changes represent increased mitochondrial biogenesis to meet increased energy demands needed to sustain PV-IN firing to maintain circuit homeostasis, particularly in the setting of emerging synaptic defects in early Aβ pathology.

### Evidence for decreased mTOR-C1 signaling in PV-INs in early A**β** pathology

The elevated abundances of several mitochondrial and metabolic proteins and decrease in synapto-dendritic proteins raise the possibility that upstream signaling pathways may be dysregulated in PV-INs. Metabolic signaling pathways such as Akt/mTOR are important regulators of mitochondrial biogenesis and turnover by impacting fusion and fission, as well as synaptic function^68–71^. Signaling via several MAPKs including ERK and p38 MAPK impact cell proliferation, synaptic function and survival. These broad pathways may therefore collectively regulate metabolism, and alteration in levels of activation in these pathways may indicate adaptive or dysfunctional responses to stress in PV-INs. We found that PV-CIBOP labeled 75 proteins that belong to Akt/mTOR and MAPK (ERK, p38 MAPK and Jnk) signaling pathways (**Fig 7F**), including 19 Akt/mTOR signaling proteins (e.g. Mtor, Rptor, Eif4b) and 49 MAPK signaling proteins (e.g. Map2k1, Ras proteins, Pak2, Akt3, Mapk3, Mapk10). Of these, few proteins showed increased levels (Rhoa, Prkca, Hras, Cacna2d1) and decreased levels (Eif4b, Mapt, Rheb) in 5xFAD PV-INs as compared to WT PV-INs (**Fig 7G**). Interestingly, these differential effects of Aβ pathology on levels of these signaling proteins were only observed in PV-IN proteomes, and not in the bulk brain proteome with the exception of Rheb, indicating the specificity of these alterations in PV-INs in early AD pathology (**Fig 7H**). Based on the ability of CIBOP to biotinylate signaling proteins in PV-INs, we performed adapted Luminex assays to detect MAPK (Erk, P38 Mapk and Jnk) and Akt/mTOR signaling phospho-proteins, specifically derived from PV neurons, as we recently described^7^. In this approach, the biotinylated phospho protein of interest is first immobilized on a bead using capture antibodies, and then their biotinylation status is detected using a streptavidin fluorophore, allowing us to directly quantify biotinylated PV-IN-derived phospho-proteins from the bulk homogenate (**Fig 7I, Supplemental Datasheet 5**). Using this approach, we found that mTOR signaling, particularly phosphorylation of mTOR and its down-stream target (p70 S6K), was decreased in 5xFAD PV-INs while MAPK pathway activation was not altered (**Fig 7J**). This pattern of decreased mTOR signaling was consistent with lower levels of Rheb (a direct activator of mTOR-C1 function), higher levels of RhoA (a known inhibitor of Rheb function) and lower levels of Eif4b (involved in translation initiation) in 5xFAD PV-INs (**Fig 7K**)^72^. Collectively, our MS PV-IN CIBOP studies and adapted Luminex analyses are indicative of decreased mTOR-C1 activity in 5xFAD PV-INs.

To assess whether this molecular signature of decreased mTOR-C1 signaling in 5xFAD PV-INs is functionally relevant, we assessed three composite measures of mTOR-C1 signaling, including autophagy (mTOR signaling inhibits autophagy)^71^, translational efficiency and protein degradation (mTOR increases translational efficiency and decreases protein degradation)^73, 74^ and synaptic plasticity (mTOR positively regulates synaptic plasticity) (**Fig 7K**)^70^. 23 GO annotated positive regulators of autophagy (GO 0010508) proteins were labeled in PV-INs and collectively, this group of proteins showed an increased abundance in 5xFAD PV-INs (**Fig 7L**). We also measured levels of LC3 (II and I) by Western blot analyses of biotinylated proteins as well as bulk brain samples from WT and 5xFAD PV-CIBOP mice and found increased LC3-II (relative of LC3-I) in 5xFAD PV-INs as compared to WT PV-INs, consistent with increased autophagy in 5xFAD PV-INs (Fig 7M). Furthermore, this difference was not apparent at the bulk brain level. Interestingly, all PV-IN (WT or 5xFAD) systematically showed higher levels of LC3-II as compared to bulk brain samples (**Fig 7M**). These patterns suggest that PV-INs, as a cell type, exhibit higher levels of autophagy as compared to the rest of the brain^5^, and this phenotype is exaggerated in early AD pathology.

Decreased mTOR-C1 signaling should also lead to decreased translational efficiency and increased protein degradation. A molecular footprint of protein turnover (resulting from protein synthesis and degradation) can be ascertained by assessing the relative abundance of long-lived and short-lived proteins in the neuronal proteome. Using a reference dataset of protein half-life estimates derived from *in vivo* isotopic labeling studies in adult mice^75^, we found that DEPs that were increased in 5xFAD PV-INs were biased towards longer-lived proteins, as compared to non-DEPs as well as compared to DEPs with decreased levels in 5xFAD PV-Ins (**Fig 7N, Supplemental Datasheet 6**). This pattern of relatively increased abundance of longer lived proteins in 5xFAD PV-INs is consistent with decreased translational efficiency and/or increased protein degradation in 5xFAD PV-INs. Lastly, we analyzed 102 proteins involved in regulation of synaptic plasticity (GO 0048167) as being functionally indicative of synaptic plasticity which is positively regulated by mTOR-C1 signaling. As a group, synaptic plasticity related proteins were decreased in 5xFAD PV-INs (**Fig 7O**), consistent with observed synaptic protein changes and physiological defects presented earlier (**Fig 5, 6**). These complementary analyses of distinct facets of the PV-IN proteome provide congruent lines of evidence for decreased mTOR signaling at various levels of the signaling axis (upstream and downstream of mTOR-C1 complex function), that are associated with increased autophagic flux, decreased translational efficiency and decreased synaptic plasticity in PV-INs at early stages of Aβ pathology.

## DISCUSSION

In this study we describe a versatile intersectional method^7, 9, 15^ allowing quantitative *in vivo* neuron-type-specific proteomics. We leveraged this approach to isolate the native-state PV-IN proteome, and were also able to examine changes in this cell type in an early-stage mouse model of Aβ pathology. In principle, our approaches should be readily adaptable to other neuron types, due to the recent expansion of novel cis-element-directed AAV approaches^76, 77^ with unparalleled cell-type-specificity. A secondary advantage to our workflow is the ability to seamlessly translate the technique across different mouse models of disease, without the need for overly complex and expensive cross-breeding strategies. Comprehensive molecular characterization of neuronal subtypes can provide critical insights into the function and modifiable pathogenic mechanisms of neurological diseases. Ideally, this should be performed at the proteomic level, while retaining the native state of neuronal proteome (i.e., protein samples from intact soma, dendrites, and axons). Current transcriptomic studies of neurons have relied on isolation of intact neurons or neuronal nuclei for RNAseq studies. The overall level of concordance between mRNA and functionally-relevant protein in the brain, is modest at best^2–4^. Further, among various cell types in the brain, neuronal gene products also exhibit the highest level of protein-mRNA discordance^78^. Proteomic studies of neurons and neuronal subtypes in the *in vivo* context have lagged behind transcriptomic advances due to several technical barriers. However recent advances in cell type-specific *in vivo* proteomic labeling approaches, such as CIBOP with TurboID, and BONCAT using mutant MetRS^79–81^, provide exciting opportunities to investigate neuronal subtype-specific molecular signatures and disease mechanisms in mouse models. For the CIBOP approach described in this study, proximity based biotinylation of proteins in the cytosol and several cellular compartments (e.g., synaptic boutons or postsynaptic densities) can be achieved by TurboID expression selectively in brain cell types of interest, by driving Cre recombinase expression via transgenic or AAV approaches, in the Rosa26^TurboID^ mouse line^7^.

CIBOP with TurboID mice has thus far been successfully applied in specific brain cell types by crossing with cell-selective Cre mouse lines (Camk2a-Cre for excitatory neurons and Aldh1l1-Cre for astrocytes) and is thus well positioned to be extended to other neuronal subtypes and glial cells^7, 81, 82^. Universal extension of cell-specific CIBOP across mouse models of disease would provide unparalleled resolution to pathological mechanisms. However, the need for further time-consuming and expensive cross-breeding represents a significant barrier to entry. Thus, here we developed an enhancer-AAV method to deploy CIBOP more rapidly in PV-INs of both WT and disease model mice. Further development of intersectional AAV approaches should also allow CIBOP to be expanded in other model species as well as to cell types outside the brain^83^.

Among neuronal subtypes, PV-INs represent a unique class of inhibitory interneurons with fast-firing properties and high metabolic activity^12, 84^. Selective dysfunction of PV-INs contributes to a variety of neurological insults, including in neurodegenerative diseases such as AD, neurodevelopmental disorders, and catastrophic early-life epilepsy^85–87^. We applied CIBOP to PV-INs throughout the forebrain by subcloning a Cre-expressing cassette into a PV-specific enhancer-AAV^9^ (AAV.E2.Cre) delivered systemically using the PHP.eB serotype, which readily crosses the BBB^15^. Delivery of AAV.E2.Cre in Rosa26^TurboID^ mice lead to PV-specific proteomic labeling via biotinylation. When coupled to MS analysis of affinity-purified biotinylated proteins, we could capture the PV-IN-specific proteome while retaining the native functional state of the cells, obviating the need for potentially detrimental cell isolation procedures. Importantly, PV CIBOP approach did not impact electrophysiological properties or density of PV-INs in the mouse brain, verifying that our CIBOP approach were representative of native-state proteomes of physiologically unperturbed PV-INs. Taken together, the PV-CIBOP approach identified unique proteomic signatures of PV-INs complementing existing transcriptomic data, serving as an important resource to the neuroscience research community.

The PV-IN proteomic signature included hundreds of proteins that were either exclusive to, or highly-enriched in PV-INs over the bulk brain proteome, which included several canonical PV-IN markers (e.g. Kv3.1-3.3, Erbb4, Ank1, Syt2)^11^. Of 294 proteins that were exclusive or highly enriched in the PV-IN over the bulk proteome, majority were not previously reported as PV-IN-specific markers based on existing PV-IN scRNAseq databases (eg. Tnpo3, Htt, Synrg, Cplx3, Mtor, Gria1). We also observed an overall modest concordance between the proteome and scRNAseq transcriptome of PV-INs, based on an analysis of over 1,800 protein-mRNA pairs in our study. PV-CIBOP labeled proteins with predicted subcellular targets in the cell body, axons and dendrites of PV-INs, including within both pre and post-synaptic compartments. Overall, the PV-IN proteome was suggestive of high metabolic, translational and mitochondrial activity, consistent with the fast-firing and rapid neurotransmission^14^ properties of these cells.

Our WT PV-CIBOP proteome was highly enriched in several proteins representing associations to neurodegenerative disease risk, including AD through altered APP cleavage (BIN1)^88^, pure tauopathies (MAPT)^89^, and synucleinopathies (SNCA, SNCB)^90, 91^. Several lines of evidence now suggest that interneuron dysfunction leads to altered circuit excitability in early stage models of APP/Aβ pathology^92, 93^. Stx1b and Gat1, among others, are examples of proteins enriched in the PV-IN dataset linked to both epileptiform activity and AD^94^. Furthermore, other neurodevelopmental and autism implicated molecules (Shank2, Syngap1, ErbB4)^95–99^ were also highly enriched in PV-INs. When contrasted to CIBOP-derived proteomes from the (predominantly pyramidal neuron) Camk2a-Cre mice, PV-IN proteomes exhibited clearly higher levels of ribosomal, endocytic, Akt/mTOR signaling, synaptic cytoskeletal, endocytic and synaptic vesicle-related proteins. In contrast, Camk2a-CIBOP signatures unsurprising were enriched in glutamatergic signaling proteins consistent with the predominantly excitatory nature of these cells. The proteomic contrast between PV-IN and Camk2a neurons in mice, integrated with recently-identified proteomic correlates of cognitive resilience in humans, revealed a disproportionate enrichment of pro-resilience proteins in the PV-IN proteome, suggesting a link between cognitive resilience and PV-INs.

To look for evidence of PV-IN-associated vulnerability in human AD, we assessed existing human post-mortem bulk brain proteomic data from controls and AD cases^48^. These post-mortem datasets are useful as they include asymptomatic AD patients (i.e., those exhibiting post-mortem neuropathological features of AD without cognitive dysfunction prior to death). We identified a module of co-expressed proteins (module M33) that was uniquely enriched in PV-IN markers (eg. Pvalb, Kcnc2, and was distinct from modules enriched in excitatory neuronal (pyramidal cell) markers. The overall level of abundance of the PV-IN proteomic module was lower in AD cases compared to both asymptomatic AD and control patients. A decrease in M33 module-associated proteins was also strongly correlated with AD neuropathological features (Aβ pathology, neurofibrillary tangles), severity of cognitive dysfunction, and rate of cognitive decline. After accounting for neuropathological hallmarks of neurodegenerative diseases, M33 still remained associated with cognitive resilience, raising the possibility that the relationship between PV-INs and cognitive resilience are independent of the type of neurodegenerative pathology. We also used complementary analytical approaches to estimate abundances of different neuronal classes, and found that PV-IN cell type estimates were among the most strongly associated with cognitive resiliency. Together, our results suggest that preservation of PV-IN function in the brain may be generally protective in AD. Although derived from bulk brain proteomes, these human findings associate PV-INs with cognitive resilience and vulnerability with AD.

Since human post-mortem brain studies cannot directly capture longitudinal impacts of aging and disease progression, we next analyzed mouse bulk brain proteomes from WT mice and 5xFAD mice which exhibit accelerated Aβ pathology, spanning a wide 1.8-14 month age range. We found that PV-enriched proteins Pvalb and Kcnc showed age-dependent increases in expression. Interestingly, Pvalb and Kcnc proteins, but not excitatory neuron-enriched Cam2ka, Slc17a7, and Vglut1, showed progressive changes in 5xFAD mice starting 3 months of age. Albeit a snapshot from bulk tissue, this suggests that of PV-IN protein levels, but not other neuronal markers, may change in early stages of APP/Aβ pathology. Somatostatin (Sst), a protein primarily expressed in cortical dendrite-targeting (non-PV fast spiking)^100, 101^ inhibitory interneurons, was similarly reduced starting at 3 months of age. As both Pvalb and Sst expression are linked to the level of circuit activity^102, 103^ these changes may reflect a differential dysregulation of interneuron activity levels at a stage where substantial plaque formations are just arising in 5xFAD mice. At the histological level, no measurable differences in PV-IN density were observed between 3 month old wild-type and 5xFAD mice, arguing against early overall cell loss of PV-INs at this early stage, but rather suggesting changes to their proteomic profile.

To evaluate PV-IN proteomic changes in response to early Aβ pathology, we compared PV-CIBOP-derived proteomes from 3 month old Rosa26^TurboID^ WT and 5xFAD mice. Over 450 DEPs were found. Proteins involved in mitochondrial function, cholesterol biosynthesis (e.g., Dhcr7), and metabolism were generally increased in PV-INs. In contrast, cytoskeletal, structural, and synapse-associated proteins were generally decreased in PV-INs. Surprisingly, alterations found in the PV-IN proteome were almost completely non-overlapping with those changes resolved from bulk brain. Since the majority of intra-neuronal APP/Aβ was detected in non-PV-INs at this stage of pathology, the observed changes in PV-INs is most likely due to Aβ from other neurons rather than due to dysfunctional Aβ processing in PV-INs. Based on these specific effects of Aβ pathology on PV-INs but not other brain cell types, extending CIBOP to other interneuron and excitatory neuron subclasses, and capturing the effects of brain region and temporality (age) in future studies will be necessary to resolve whether PV-IN protein levels are profoundly affected in early AD models, or rather, are part of a continuum of emerging cell type autonomous alterations across different brain regions.

Initial PV-CIBOP studies in WT mice found substantial enrichment of AD-associated proteins identified in a genome-wide risk analysis (MAGMA AD) study, as well as enrichement of pro-resilience proteins in the PV-IN proteome in contrast to Camk2a neurons^34, 35, 48^. Therefore, we asked whether MAGMA AD proteins would also be disrupted in our early AD model PV proteome, and indeed, cross-referenced DEPs in 5xFAD matched with 20 MAGMA AD genes. Furthermore, several proteins associated with cognitive resilience were reduced in 5xFAD PV cells. Of these MAGMA AD and pro-resilience related proteins, we identified several with shared functional involvement with presynaptic vesicle fusion/exocytosis/release (Cplx1, Cplx2, Stx1b, Elfn1, Rab3c, Rims1)^35^. To examine whether this signature of synaptic dysfunction was expressed functionally, we used PV-IN-specific optogenetic approaches in two independent models of early APP/Aβ pathology. At cortical PV-to-pyramidal synapses, both studies clearly point to disturbances in presynaptic function. In particular, changes in vesicular release probability appears likely. Additional work is necessary to characterize whether changes in Cplx1, Cplx2, Stx1b, Elfn1, Rab3c, or Rims1, or yet other mechanisms can explain our findings. Several studies have also shown an emergence of inhibitory postsynaptic dysfunction across a number of APP/Aβ models^104–107^. Further longitudinal evaluation of PV-IN and other inhibitory cell synaptic mechanisms are warranted. Nonetheless, the synaptic mechanisms identified at PV synapses in this study may represent opportunities for early therapeutic intervention. Deployment of combined cell-type-specific CIBOP and functional evaluations with high vulnerability in early AD represent a logical next step.

Despite minimal plaque burden in 3 month old 5xFAD mice^21^, the significant shifts in the 5xFAD PV-IN proteome may represent a homeostatic response to prior changes in neuronal and circuit behavior and organization^65, 108, 109^ known to occur young, pre-plaque APP/Aβ models, including in younger (< 3 month old) 5xFAD mice^16, 93, 110, 111^. Relatedly, a signature of circuit instability is also present in human patients with mild cognitive impairment and AD^19, 112, 113^. To compensate for this early circuit dysfunction, PV-INs are well suited to homeostatically respond^114^, but this process could induce significant metabolic stress and impose a higher metabolic demand to sustain this compensation. Indeed, mitochondrial impairments have been observed prior to extensive pathology in APP/Aβ model mice^115, 116^. In our PV-CIBOP proteomes, we found an extensive signature of stress-responsive proteins (Armt1, Rhob, Gstm1, RhoA, Tmco1, Akr1b3, Gcn1, Hras, Cul3, Pdk2, Rap2a, Flot1) in 5xFAD as compared to WT. Of note, RhoA activation increases Aβ and tau pathology and co-localizes with NFTs in human brain^117, 118^. In contrast to the overall synaptic effects of early AD pathology in PV-INs, we observed a marked increase in mitochondrial and metabolic proteins in PV-INs. This increase could be reflective of a protective or compensatory responses (via increased mitochondrial biogenesis to sustain higher metabolic demand). Other compensatory signatures observed in 5xFAD PV-INs included increased Dhcr7 for de-novo cholesterol biosynthesis in neurons, increased Apeh to process Aβ oligomers along with increased autophagy as supported by increased levels of positive regulators of autophagy and increased lapidated form of LC3 (LC3 II). Conversely, a detrimental/dysfunctional response (e.g., accumulation of dysfunction mitochondria) is also possible. We noted that mitochondrial functional proteins and Complex I, III, IV, V proteins were selectively increased in 5xFAD PV-INs while a smaller group of mitochondrial structural, dogma, and Complex II proteins were not. Therefore, follow-up studies focusing on mitochondrial structure and function specifically in PV-INs are warranted to better understand the basis and consequences of these mitochondrial alterations.

Among the proteins most highly increased in 5xFAD PV-INs, we validated the Complex IV protein Cox5a via western blotting and MS approaches. In bulk tissue, Cox5a protein level was minimally decreased at 3 months of age although age-dependent decline in Cox5a was observed at later timepoints. In contrast, Cox5a protein was increased in PV-INs, further supporting the notion that increased mitochondrial respiratory chain proteins may represent a unique compensatory mechanism in PV-INs that is not occurring in the rest of the brain. Cox5a and many other respiratory chain proteins that were increased in 5xFAD PV-INs also demonstrated very high level of mRNA/protein discordance, whereby PV-INs express high levels of these transcripts but very low levels of proteins under non-disease conditions. It is therefore possible that early 5xFAD pathology impacts post-transcriptional and translational regulation of these mitochondrial genes. Future transcriptomics studies of PV-INs in AD models may help address this possibility. Taken together, the molecular phenotype of 5xFAD PV-INs is indicative of a significant cellular stress response occurring in 3 month old PV-INs, comprising both compensatory and maladaptive events, which is not evident in the bulk proteome at this age. Furthermore, we present several lines of evidence from bulk brain and PV-IN-specific experiments, and human brain proteomic analyses, that PV-IN proteomic signatures and cognitive resiliency are linked. Therefore, understanding the mechanisms for this compensation could provide therapeutic insights for future studies.

Metabolic shifts and mitochondrial biosynthesis are regulated by signaling pathways such as Akt/mTOR and MAPK^68–71^. We also observed high levels of proteins involved in both Akt/mTOR (e.g. Mtor, Eif4b) and MAPK (e.g. Erk and Mek proteins) signaling pathways in PV INs. Therefore, we hypothesized that mTOR signaling may be altered in 5xFAD PV-INs. We directly measured biotinylated phospho-proteins indicative of levels of activity of these pathways specifically in PV-INs by leveraging an adapted Luminex immunoassay method recently validated for CIBOP-based studies. Using both MS-based quantification of signaling proteins and Luminex-based assays of phospho-proteins, we found evidence of decreased mTOR (mTOR-C1) signaling in PV-INs but not in bulk brain tissue. Further analyses of our MS data and follow-up validation studies indicate functional consequences of decreased mTOR-C1 signaling in PV-INs, including augmented autophagic flux, decreased translational efficiency or increased protein degradation, as well as synaptic dysfunction. We also observed a general pattern of increased autophagic flux in PV-INs as a cell type, as compared to bulk brain tissue. Taken together, we provide evidence for early dysregulation of mTOR signaling in PV-INs as a potential upstream mechanism for mitochondrial and metabolic alterations as well as synaptic dysfunction occurring selectively in PV-INs in early stages of AD pathology in 5xFAD mice.

Limitations of our work relate to technical considerations of both AAV-based PV-IN targeting, and potential proteomic biases of the CIBOP approach. Currently, CIBOP leads to cell type-specific expression of TurboID-NES, which contains a nuclear export sequence for preferential proteomic labeling outside the nucleus. This may bias the PV-IN proteome away from nuclear proteins as well as from proteins present within the lumen of organelles (e.g. ER/Golgi, mitochondria, lysosomes)^7, 82^. Whether removal of the NES impacts the nature of the PV-IN proteome, remains to be determined Another consideration is that our AAV.E2.Cre strategy targets PV-INs as a whole, although several PV-IN subtypes have been identified by scRNAseq studies^1^ (i.e., chandelier cells and several basket cell PV types). It is therefore possible that the proteomic signatures of these different PV-IN subtypes are non-uniform. Thus, our initial PV-CIBOP derived proteome may not accurately describe the proteomic granularity which may further exist within PV-IN interneuron classes. Further studies with increased PV subtype-specificity or physiological and morphological studies using CRISPR or related methods to examine individual proteins may be useful in this regard.

In summary, our integrative PV-CIBOP approach revealed a novel native-state proteomic signature for a single, highly important interneuron class in the mouse brain. Comparison of PV-CIBOP proteomic signatures with human post-mortem data suggests selective synaptic and metabolic PV-IN vulnerabilities in early AD pathogenesis. These findings provide a strong rationale to investigate early proteomic changes occurring in PV and other inhibitory neuron types in mouse models of AD pathogenesis, using an analogous enhancer AAV CIBOP approaches.

## METHODS

### Reagents

A detailed list of reagents used in these studies, including antibodies, is provided in Supplemental Datasheet 7.

### Animals

C57BL/6J (wildtype[WT]) mice (JAX #000664), Rosa26-TurboID (C57BL/6-Gt(ROSA)26Sortm1(birA)Srgj/J, JAX #037890)^7^, Camk2a-Cre-ert2 (B6;129S6-Tg(Camk2a-cre/ERT2)1Aibs/J, JAX #012362)^119^ and 5xFAD (B6.Cg-Tg(APPSwFlLon,PSEN1*M146L*L286V)6799Vas/Mmjax, JAX #034848)^17, 21^ mouse lines were used for experiments in this study and genotyping was performed using primers and polymerase chain reaction (PCR) conditions listed on the vendor website (Jackson labs). All animals were maintained on the C57BL6/L background, following at least 10 serial backcrosses if originally derived from a different or mixed background. Male and female mice were used for all experiments with data collected from ≥ 3 mice per experimental condition for all experiments. Animals were housed in the Department of Animal Resources at Emory University under a 12 h light/12 h dark cycle with ad libitum access to food and water and kept under environmentally controlled and pathogen-free conditions. All experiments involving animal procedures were approved by the Emory University Institutional Animal Care and Use Committee (IACUC, PROTO201700821) and were in accordance with the ARRIVE guidelines.

### Retro-Orbital AAV injections

The same AAV retro-orbital injection was given to male and female mice of each genotype as previously described^15^. Rosa26^TurboID/wt^ and WT control mice were briefly (∼2 minutes) anesthetized with 1.5-2% isoflurane to perform the injection. AAV(PHP.eB).E2.Cre.2A.GFP virus (2.4E+11 vector genomes) and AAV(PHP.eB).Flex.Tdtom (3.15E+11 vector genomes) were co-injected (final volume; 65μl in sterile saline) to target and label PV interneurons throughout the cortex. Injections into the retro-orbital sinus of the left eye were performed with a 31G x 5/16 TW (0.25 mm) needle using an insulin syringe. Fresh syringes were used for each mouse. Mice were kept on a heating pad for the duration of the procedure until recovery (< 5 minutes) and then returned to their home cage. After 3 weeks post-injection, mice were provided with biotin water continuously. Biotin water was administrated for 2 weeks until acute slice sample collection (total of 5 weeks post-RO injection).

### Tissue collection

Mice were fully anesthetized with isoflurane and euthanized by decapitation. Mice brains were immediately removed by dissection. The right hemisphere of the brain was snap frozen on dry ice either intact or after regional dissection (cerebellum, brain stem, cortex, hippocampus, and striatum/thalamus) for downstream proteomics analysis while left hemisphere were used for electrophysiological recording and immunostaining.

### Acute Slice Preparation

Brains were immediately removed by dissection in ice-cold cutting solution (in mM) 87 NaCl, 25 NaHO3, 2.5 KCl, 1.25 NaH2PO4, 7 MgCl2, 0.5 CaCl2, 10 glucose, and 7 sucrose. The left hemisphere of the brain was used for electrophysiological recording and immunostaining. For electrophysiology, brain slices (300 μm) were immediately prepared in the coronal or sagittal plane using a vibrating blade microtome (VT1200S, Leica Biosystems) in the same solution. Slices were transferred to an incubation chamber and maintained at 34°C for ∼30 min and then 23-24°C thereafter for patch clamp recordings. Solutions were equilibrated and maintained with carbogen gas (95% O2/5% CO2) throughout the entire day. Remaining tissue from the left hemisphere was transferred to 4% paraformaldehyde (PFA) in 0.1 M phosphate-buffered saline (PBS)PFA in PBS at 4°C for immuno-histological analysis.

### CIBOP studies

PV-CIBOP studies in WT mice were performed by single retro-orbital injections of AAV (AAV(PHP.eB).E2.Cre.2A.GFP) to Rosa26^TurboID/wt^ mice. Rosa26^TurboID/wt^ (PV-CIBOP) were also crossed with 5xFAD (hemizygous) to derive 5xFAD (hemi)/Rosa26^TurboID/wt^ (5xFAD/PV-CIBOP) and littermate WT/PV-CIBOP) animals. Camk2a-CIBOP experiments were performed in Camk2a-Cre-ert2^het^/Rosa26^TurboID/wt^ mice. Tamoxifen was injected (intraperitoneally or i.p., 75mg/kg/dose in corn oil x for 5 consecutive days), followed by 3 weeks to allow Cre-mediated recombination and TurboID expression after which biotinylation (37.5 mg/L in drinking water) was performed for 2 weeks^7^. In the case of PV-CIBOP (on WT or 5xFAD backgrounds), AAV injections were followed by 3 weeks to allow recombination after which biotinylation was performed as above. For all CIBOP studies, control animals included Cre-only (in the case of Camk2a-CIBOP experiments) or Rosa26^TurboID/wt^ mice (for PV-CIBOP experiments). Control groups in PV-CIBOP studies also received AAV E2.Cre injections for fair comparisons. In Camk2a-CIBOP studies, all experimental groups received tamoxifen to account for tamoxifen mediated effects. No tamoxifen was needed for PV-CIBOP experiments (non-inducible Cre). Recombination period (after inducing Cre via tamoxifen or delivery of AAV E2.Cre) and biotinylation period after recombination were kept constant across all studies. CIBOP studies were completed at 12 to 13 weeks of age.

### Electrophysiology

For whole-cell current clamp recordings, acute slices were continuously perfused in a recording chamber (Warner Instruments) with (in mM) 128 NaCl, 26.2 NaHO3, 2.5 KCl, 1 NaH2PO4, 1.5 CaCl2, 1.5 MgCl2 and 11 glucose, maintained at 30.0±0.5°C. The recording solutions were equilibrated and maintained with carbogen gas (95% O2/5% CO2) in a closed loop system for the entire experiment. PV neurons were targeted for whole-cell recordings in layer 5 somatosensory cortex as previously described using combined gradient-contrast video microscopy and epifluorescent illumination either custom-built or commercial (Olympus) upright microscopes^16^. Although our AAV expressed a GFP downstream of a 2A linker (**Fig 1**), GFP fluorescence was too dim, hence co-injection with a floxed tdTom for fluorescence-targeted patch-clamp of PV interneurons. Recordings were obtained using Multiclamp 700B amplifiers (Molecular Devices). Signals were low pass filtered at 4–10 kHz and sampled at 50 kHz with the Digidata 1440B digitizer (Molecular Devices). Borosilicate patch pipettes (World Precision Instruments) were filled with an intracellular solution containing (in mM) 124 potassium gluconate, 2 KCl, 9 HEPES, 4 MgCl2, 4 NaATP, 3 l-ascorbic acid, and 0.5 NaGTP. Pipette capacitance was neutralized in all recordings and electrode series resistance compensated using bridge balance in current clamp. Liquid junction potentials were uncorrected.

For PV interneuron current clamp experiments, membrane potentials were maintained at –70 mV using a constant current bias. AP trains were then initiated by somatic current injection (300 ms) normalized to the cellular capacitance in each recording measured immediately in voltage clamp after breakthrough^16, 120^. For quantification of individual AP parameters, the first AP in a spike train was analyzed for all cells. Passive properties were determined by averaging the responses of several 100ms long, -20 pA steps during each recording.

For voltage clamp recordings, cells were filled with an intracellular solution containing (in mM) 120 CsMeSO4, 10 HEPES, 5 TEA.Cl, 4 Na2ATP, 0.5 Na2GTP, 2 MgCl2, 10 L-Ascorbic Acid, and 3 Qx314. Spontaneous excitatory and inhibitory postsynaptic currents (sEPSCs and sIPSCs) were recorded at a holding voltage of -70 and 0 mV interleaved for one second each for 3-5 minutes. Signals were filtered with a Bessel 10 kHz low-pass filter and sampled at 50 kHz. All recordings had a series resistance of < 20 MΩ. Event detection was carried out using Clampfit (Molecular Devices) using a template matching algorithm and were manually curated following posthoc 4 kHz low-pass filtering. Events were compared against the template within a match threshold of 2.5. Events with a rise time of < 0.15 ms and a decay time less than double the rise time were excluded. All statistical tests were completed in Prism (GraphPad) by using unpaired t-tests for cell averages, and p < 0.05 was considered statistically significant. All group statistics are presented as cumulative frequency distribution or mean ± SEM (standard error of mean).

For optogenetic slice experiments, C1V1 was activated using an unfiltered amber LED (M590L3; Thorlabs) centered on l = 596 nm (±15 nm). ChETA was excited using blue light from an unfiltered LED (M470L3; Thorlabs) centered on l = 461 nm (± 20 nm). LEDs were rapidly modulated with time-locked TTL pulses from the electrophysiology software using short pulses with a current controller (LEDD1B; Thorlabs). Due to kinetics differences between ChETA and C1V1, 0.15-0.3 and 4ms pulses (respectively) were found to be optimal to reliably elicit PV pyramidal cell IPSCs every trial with minimal jitter. To eliminate potential sources of variation between experiments, these parameters and the amber/blue light power at the objective remained unaltered for all optogenetic control and test experiments.

### Stereotactic surgery for optogenetics

PV-IN-specific optogenetic studies were performed in separate cohorts of WT and 5xFAD mice, or in PV-Cre (Jax strain #:017320) mice for hAPP-AAV studies. For C1V1 experiments in 5xFAD (or WT controls), AAV(PHP.eB)-E2-C1V1-eYFP (addgene 135633) was co-injected with AAV1.CamKII(1.3).eYFP.WPRE.hGH (addgene 105622) at a 1:1 ratio. For ChETA experiments, AAV1-Ef1a-DIO-ChETA-EYFP (addgene 26968) was co-injected with an hAPP-expressing virus AAV(PHP.eB).EF1a.hAPP.oPRE (hAPP RefSeq NM_000484.4) or an equivalent amount of sterile saline in controls. Injections were all performed in the SBFI region of S1 cortex. For injections, mice were head-fixed in a stereotactic platform under continuous isoflurane anesthesia (1.8–2.0%). Thermoregulation was provided by a heat plate with a rectal thermocouple for biofeedback, to maintain core body temperature near 37°C. A small incision was made and a craniotomy cut in the skull (<0.5 μm in diameter) to allow access for a glass microinjection pipette. Coordinates (in mm from Bregma) for microinjection were X = ±3.10– 3.50; Y = −2.1; α = 0°; Z = 0.85–0.95. Viral solutions (stock titers were ∼1.0 × 10^13^ vg/mL, except for AAV.EF1a.hAPP, which was 1.0 x 10^12^ vg/ml) were injected slowly (L0.02 μL min^-1^) using Picospritzer-directed short pulses (∼0.3 μL total). After ejection of virus, the micropipette was held in place (∼5 min) before withdrawal. The scalp was closed first with surgical sutures and Vetbond (3M) tissue adhesive thereafter and the animal was allowed to recover under analgesia (carprofen and buprenorphine SR). After allowing for onset of expression (1 or 3 weeks for C1V1/YFP or ChETA/hAPP, respectively), animals were sacrificed, and acute slices harvested for patch clamp studies as detailed above.

### Immunohistochemistry (IHC) and immunofluorescence microscopy

IHC was performed on brain slices obtained from CIBOP studies. Tissue was transferred immediately after euthanizing the animals in 4% PFA or after completion of electrophysiological recordings. Brain tissue/slices were post-fixed in 4% PFA in PBS at 4°C overnight then subsequently transferred into 30% sucrose solution until cryo-sectioning. After embedding in optimal cutting temperature (OCT) compound, the slices were further cut coronally or sagittally into 40-μm-thick sections on a cryostat (Leica Biosystems). IHC and immunofluorescence protocols used were as previously published^7^. The tissue sections were transferred into the cryoprotectant or directly mounted on the charged glass slides and stored at -20°C until use. For immunofluorescence (IF) staining, 40Lµm thick free-floating brain sections or glass slides were washed, blocked and permeabilized by incubating in TBS containing 0.30% Triton X-100 and 5% horse serum for 1 h at room temperature. If desired, then the antigen retrieval was performed in Tris-EDTA buffer (10 mM Tris base, 1mM EDTA, 0.05% tween-20) pH9 for 30 minutes at 65°C before the blocking and permeabilization step. Primary antibodies were diluted in TBS containing 0.30% Triton TX-100 and 1% horse serum. After overnight incubation at 4°C with primary antibodies, the sections were rinsed 3x in TBS containing 1% horse serum at room temperature for 10 min each. Then, the sections were incubated in the appropriate fluorophore-conjugated secondary antibody at room temperature for 2 h in the dark. The sections were rinsed once and incubated with DAPI (1µg/ml) for 5 min, washed 3x in TBS for 10 min, dried, and cover slipped with ProLong Diamond Antifade Mountant (P36965; ThermoFisher,). All the primary and secondary antibody detail including dilution used are listed in **Supplemental Datasheet 7**. We optimized several existing antibodies to detect Pvalb protein by IHC (**Supplemental Datasheet 7**). Since the AAV E2.Cre.GFP labels all PV-INs with very high efficiency, cells expressing GFP were used as the reference standard for PV-INs and Pvalb antibodies with highest levels of agreement with GFP-positive PV-INs were used interchangeably for IHC studies, to allow species compatibility of primary antibodies. In addition, the guinea pig Pvalb antibody (195004; Synaptic Systems) preferentially labeled the synapto dendritic compartment of PV-INs. IHC studies were also performed on experimental animals from CIBOP studies, as well as from non-CIBOP WT and 5xFAD mice using sagittal sections of entire hemispheres.

Images of the same region across all samples were captured as z-stacks using the Keyence BZ-X810, except for images in **Figure S5** (Parvalbumin IHC quantification). Some of the z stacked images of entire brain were stitched together to allow regional comparison based on level of biotinylation. Images for quantification of Parvalbumin staining were obtained with a two photon laser scanning microscope (2pLSM) using a commercial scan head (Ultima; Bruker Corp) fitted with galvanometer mirrors (Cambridge Technology) using a 60x, 1.0 NA objective. Parvalbumin levels were quantified in an analogous fashion to that described previously^121^, but at higher magnification to resolve potential differences across different cortical layers. All image processing was performed either using the Keyence BZ-X810 Analyzer or Image J software (FIJI Version 1.53).

### Tissue processing for protein based analysis, including Western Blot (WB)

Tissue processing for proteomic studies, including MS studies, were performed in a previous CIBOP study^7^. Frozen brain tissues (whole brain homogenate excluding the cerebellum for studies in Fig 1 and whole cortical samples for studies in Figs 2 and 5) either intact or dissected cortex, was weighed and added to 1.5 mL Rino tubes (Next Advance) containing stainless steel beads (0.9–2 mm in diameter) and six volumes of the tissue weight in urea lysis buffer (8 M urea, 10 mM Tris, 100LmM NaH2PO4, pH 8.5) containing 1X HALT protease inhibitor cocktail without EDTA (78425, ThermoFisher). Tissues were homogenized in a Bullet Blender (Next Advance) twice for 5 min cycles at 4 °C. Tissue were further sonicated consisting of 5Lseconds of active sonication at 20% amplitude with 5Lseconds incubation periods on ice. Homogenates were let sit for 10 minutes on ice and then centrifuged for 5 min at 12,000 RPM and the supernatants were transferred to a new tube. Protein concentration was determined by BCA assay using Pierce™ BCA Protein Assay Kit (23225, Thermofisher scientific). For WB analyses, 10µg of protein from brain lysates were used to verify TurboID expression (anti-V5) and biotinylation (streptavidin fluorophore conjugate). Standard WB protocols, as previously published, were followed^7^.

Other proteins detected by WB also included beta actin, Pvalb and LC3 (see **Supplemental Datasheet 7** for antibodies/dilutions). All blots were imaged using Odyssey Infrared Imaging System (LI-COR Biosciences) or by ChemiDoc Imaging System (Bio-Rad) and densitometry was performed using ImageJ software.

### Enrichment of biotinylated proteins from CIBOP brain

As per CIBOP protocols previously optimized by our group^7^, biotinylated proteins were captured by streptavidin magnetic beads (88817; Thermofisher Scientific) in 1.5 mL Eppendoft LoBind tubes using 83uL beads per 1mg of protein in a 500 µL RIPA lysis buffer (RLB)(50 mM Tris, 150 mM NaCl, 0.1% SDS, 0.5% sodium deoxycholate, 1% Triton X-100). In brief, the beads were washed twice with 1 ml of RLB and 1 mg of protein were incubated in 500 µl of total RPL. After incubation at 4 deg C for 1 h with rotation, beads were serially washed at room temperature (twice with 1 mL RIPA lysis buffer for 8 min, once with 1LmL 1LM KCl for 8 min, once with 1LmL 0.1LM sodium carbonate (Na2CO3) for ∼10 s, once with 1LmL 2LM urea in 10 mM Tris-HCl (pH 8.0) for ∼10 s, and twice with 1 mL RIPA lysis buffer for 8 min), followed by 1 RIPA lysis buffer wash 4 final PBS washes. Finally, after placing the tubes on the magnetic rack, PBS was removed completely, then the beads were further diluted in 100 µl of PBS. The beads were mixed and 10% of this biotinylated protein coated beads were used for quality control studies to verify enrichment of biotinylated proteins (including WB and silver stain of proteins eluted from the beads). Elution of biotinylated protein was performed by heating the beads in 30 μL of 2X protein loading buffer (1610737; BioRad) supplemented with 2LmM biotin + 20 mM dithiothreitol (DTT) at 95 °C for 10 min. The remaining 90% of sample were stored at -20*C for western blot or mass spectrometric analysis of biotinylated protein.

### Western blotting

To confirm protein biotinylation, 10 μg of tissue lysates were resolved on a 4–12% Bris-Tris gel (M00668, GenScript) and transferred onto a nitrocellulose membrane. The membranes were washed once with TBS-T (0.1% tween-20) and then blocked with Start Block (37543, Thermofisher Scientific) and probed with streptavidin-Alexa 680 diluted in Start Block for 1 h at room temperature. After blocking, incubation with primary antibodies was performed overnight at 4°C. Further, the membranes were washed 3 times 10 minutes each and incubated with an appropriate secondary antibody conjugated with horseradish peroxidase-conjugated or another fluorophore. Proteins were detected using the enhanced chemiluminescence method (ECL) (1705060; BioRad).Validation of MS enriched protein were performed with 100 ug equivalent of protein eluted with beads. The quantification of each band was performed by densitometric measurement.

### Protein digestion, MS, protein identification and quantification

On-bead digestion of proteins (including reduction, alkylation followed by enzymatic digestion by Trypsin and Lys-C) from SA-enriched pulldown samples (1mg protein used as input) and digestion of bulk brain (input) samples (50 µg protein), were performed as previously described with no protocol alterations^7^. In brief, after removal of PBS from remaining 90% of streptavidin beads (10% used for quality control using western blot and silver stain) were resuspended in 225 ul of 50 mM ammonium bicarbonate (NH4HCO3) buffer. Biotinylated proteins were then reduced with 1 mM DTT and further alkylated with 5 mM iodoacetamide (IAA) in the dark for 30 min each on shaker. Proteins were digested overnight with 0.5 µg of lysyl (Lys-C) endopeptidase (127-06621; Wako) at RT on shaker followed by further overnight digestion with 1 µg trypsin (90058; ThermoFisher Scientific) at RT on shaker. The resulting peptide solutions were acidified to a final concentration of 1% formic acid (FA) and 0.1% triflouroacetic acid (TFA), desalted with a HLB columns (Cat#186003908; Waters). The resulting protein solution was dried in a vacuum centrifuge (SpeedVac Vacuum Concentrator). Detailed methods for this protocol have been previously published^7^. Lyophilized peptides were resuspended followed by liquid chromatography and MS (Q-Exactive Plus, Thermo, data dependent acquisition mode) as per previously published protocols^7^. MS raw data files were searched using Andromeda, integrated into MaxQuant using the mouse Uniprot 2020 database as reference (91,441 target sequences including V5-TurboID). All raw MS data as well as searched MaxQuant data before and after processing to handle missing values, will be uploaded to the ProteomeXchange Consortium via the PRIDE repository^122^. As previously published, methionine oxidation (+15.9949LDa) and protein N-terminal acetylation (+42.0106LDa) were included as variable modifications (up to 5 allowed per peptide), and cysteine was assigned as a fixed carbamidomethyl modification (+57.0215LDa). Only fully tryptic peptides with up to 2 missed cleavages were included in the database search. A precursor mass tolerance of ±20Lppm was applied prior to mass accuracy calibration and ±4.5Lppm after internal MaxQuant calibration. Other search parameters included a maximum peptide mass of 4.6 kDa, minimum peptide length of 6 residues, 0.05LDa tolerance for orbitrap and 0.6LDa tolerance for ion trap MS/MS scans. The false discovery rate (FDR) for peptide spectral matches, proteins, and site decoy fraction were 1 %. Other quantification settings were similar to prior CIBOP studies^7^. Quantitation of proteins was performed using summed peptide intensities given by MaxQuant. We used razor plus unique peptides for protein level quantitation. The MaxQuant output data were uploaded into Perseus (Version 1.6.15) and intensity values were log2 transformed, after which data were filtered so that >50% of samples in a given CIBOP group expected to contain biotinylated proteins, were non-missing values. Protein intensities from SA-enriched pulldown samples (expected to have biotinylated proteins by TurboID) were normalized to sum column intensities prior to comparisons across groups. This was done to account for any variability in level of biotinylation as a result of variable Cre-mediated recombination, TurboID expression and/or biotinylation^7^.

### Analyses of MS data and bioinformatics analyses

Within each MS study, we compared bulk proteomes to SA-enriched proteomes to confirm that expected proteins (from either PV-INs or Camk2a neurons) were indeed enriched while non neuronal proteins (e.g. glial proteins) were de-enriched as compared to bulk brain proteomes. We also identified proteins unique to bulk or SA-enriched pulldown samples. Within SA-enriched biotinylated proteins, we restricted our analyses to those proteins that were confidently biotinylated and enriched (based on statistical significance unadj. P<0.05 as well as 2-fold enrichment in biotinylated vs. non-biotinylated samples). This allowed us to exclude proteins that were non-specifically enriched by streptavidin beads. Within biotinylated proteins, group comparisons were performed using a combination of approaches, including differential abundance analysis, hierarchical clustering analysis (Broad Institute, Morpheus, https://software.broadinstitute.org/morpheus), as well as PCA, (in SPSS Ver 26.0 or R). Differential abundance analyses were performed on log2 transformed and normalized intensity values using two-tailed unpaired T-test for 2 groups assuming equal variance across groups or one-way ANOVA + post-hoc Tukey HSD tests for >2 groups). Unadjusted and FDR-corrected comparisons were performed, although we relied on unadjusted p-values along with effect size (fold-enrichment) to improve stringency of analyses. After curating lists of differentially enriched proteins, gene set enrichment analyses (GSEA) were performed (AltAnalyze Ver 2.1.4.3) using all proteins identified across bulk and pulldown proteomes as the reference (background list). Ontologies included GO, Wikipathways, KEGG, Pathway Commons, as well as prediction of upstream transcriptional and micro RNA regulators (all included in AltAnalyze Ver 2.1.4.3). Ontologies representative of a given group were selected based on enrichment scores (Fisher test p<0.05). We used SynGO to identify the types of known synaptic proteins (in pre as well as post-synaptic compartments, and different functional classes) identified in CIBOP studies^123^.Protein-protein-interactions between proteins within lists of interest were examined using STRING (https://string-db.org/cgi/input?sessionId=bqsnbjruDXP6&input_page_show_search=on)^123^.

We also performed GSVA of DEPs identified in bulk as well as PV-IN proteomes from WT and 5xFAD PV-CIBOP mice to complement GSEA^124, 125^. As previously published, statistical differences in enrichment scores for each ontology comparing two groups, were computed by comparing the true differences in means against a null distribution which was obtained by 1000 random permutations of gene labels. Benjamini & Hochberg false discovery rate adjusted p values <0.05 were considered significant. The reference gene sets for GSVA were the M5 (Mouse) Ontology Gene Sets from MSigDB (https://www.gsea-msigdb.org/gsea/msigdb/mouse/collections.jsp?targetSpeciesDB=Mouse#M5).

### Luminex immunoassay for signaling phospho-protein quantification from mouse brain

Multiplexed Luminex immunoassays were used to measure phosphoproteins in the MAPK (Millipore 48-660MAG) and PI3/Akt/mTOR pathways (Millipore 48-612MAG). The PI3/Akt/mTOR panel included pGSK3α (Ser21), pIGF1R (Tyr1135/Tyr1136), pIRS1 (Ser636), pAkt (Ser473), p-mTOR (Ser2448), p70S6K (Thr412), pIR (Tyr1162/Tyr1163), pPTEN (Ser380), pGSK3β (Ser9), pTSC2 (Ser939) and RPS6 (Ser235/Ser236). The MAPK panel detected pATF2 (Thr71), pErk (Thr185/Tyr187), pHSP27 (Ser78), pJNK (Thr183/Tyr185), p-c-Jun (Ser73), pMEK1 (Ser222), pMSK1 (Ser212), p38 (Thr180/Tyr182), p53 (Ser15) and pSTAT1 (Tyr701). We performed adapted Luminex assays as previously described^7^ to directly quantify biotinylated proteins in PV-CIBOP samples (from WT and 5xFAD CIBOP animals), whereby the biotinylated phospho-protein of interest is first immobilized on a bead using capture antibodies, and then their biotinylation status is detected using a streptavidin fluorophore (Streptavidin-PE), to directly quantify biotinylated PV-IN-derived phospho-proteins from the bulk homogenate. Luminex assays were read on a MAGPIX instrument (Luminex). As per published protocols, we performed linear ranging for every experiment and sample type prior to full assay runs^7^. Using this approach, any signal arising from non-biotinylated (non-CIBOP) control samples is the background/noise level, which was subtracted from signals derived from CIBOP animals. We also additionally normalized these background-subtracted signals based on TurboID protein levels quantified by MS, to account for any unequal biotinylation across samples. Data were analyzed with and without this TurboID normalization, and no meaningful differences were observed between approaches, therefore the TurboID-normalized data were statistically analyzed and presented in the results.

### Analysis of existing mouse brain TMT-MS data

We used a subset of the data from a larger mouse brain TMT-MS study of aging and 5xFAD disease pathology, and a complete description of this mouse TMT-MS study including expression data after batch correction are available online (https://www.synapse.org/#!Synapse:syn27023828); and data relevant to this study are included in the supplemental data (**Supplemental Datasheet 4).** Briefly, TMT-MS was performed on whole cortical brain homogenates from 43 WT and 43 5xFAD mice (ages 1.8 mo. to 14.4 months, n=8, equally balanced based on sex). Standard tissue processing and TMT-MS pipelines were used, as we have previously published^48^. Brain samples were homogenized using a bullet blender with additional sonication in 8M Urea lysis buffer containing HALT protease and phosphatase inhibitor (78425, ThermoFisher). Proteins were reduced, alkylated and digested (Lysyl endopeptidase and Trypsin), followed by peptide cleanup and TMT (16-plex kit) peptide labeling as per manufacturer’s instructions. We included one global internal standard (GIS) per TMT plex batch to facilitate normalization across batches. All samples in a given batch were randomized across six TMT batches, while maintaining nearly-equal representation of age, sex and genotype across all six batches. A complete description of the TMT mass spectrometry study, including methods for sample preparation, mass spectrometry methodology and data processing, are available online (https://www.synapse.org/#!Synapse:syn27023828). Mass spectrometry raw data were processed in Proteome Discover (Ver 2.1) and then searched against Uniprot mouse database (version 2020), and then processed downstream as described for human brain TMT mass spectrometry studies above. Batch effect was adjusted using bootstrap regression which modelled genotype, age, sex and batch, but covariance with batch only was removed^126^. From the 8,535 proteins identified in this mouse brain proteome, we analyzed data related to known markers of distinct classes of mouse neurons and glial subtypes, based on published bulk and single cell RNAseq studies, as well as markers of AD-associated pathology (including hAbeta42 peptide and Apoe).

### Analysis of human brain proteomic data, brain cell type estimates and association with neuropathological and cognitive traits

Single nucleus Allen brain atlas snRNA data was downloaded from https://cells.ucsc.edu/?ds=allen-celltypes+human-cortex+various-cortical-areas&meta=class_label and processed in R to generate a counts per million normalized reference matrix with 47,509 non-excluded cell nuclei assigned to any of 19 cell type clusters^1, 50^. The 598 sample Banner+ROSMAP consensus proteome protein profiles of bulk dorsolateral prefrontal cortex (BA-9) from postmortem human donors was the bulk brain data for deconvolution, or ultimately, across-sample, within cell type relative abundance estimation^48^. EnsDeconv was run with some adjustment per: https://randel.github.io/EnsDeconv/reference/get_params.html and https://randel.github.io/EnsDeconv/127. Briefly, the 5 marker identification methods used to get the top 50 markers by each method were t, wilcox, combined, "none" (i.e., all genes in the snRNA reference as a profile), and regression. All methods were run for both untransformed and log2-transformed data. CIBERSORT was used as the most efficient deconvolution method with a low profile for RAM use and CPU time, and estimates from 9 of 10 successful combinations of the above marker selection and transformation methods with CIBERSORT estimation . The nine individual marker selection methods produced a redundant total of 350 marker genes, and of these, genes present in all 9 of the lists for each respective cell type were kept as a consensus list of markers. These consensus lists (**Supplemental Datasheet 3**) were used as input into the GSVA R package implementation of the ssGSEA algorithm^49^. Finally, ssGSEA estimates of within-cell type relative abundances across the 488/598 samples in the published consensus protein network^48^ were correlated to the 44 module eigenproteins (MEs), which are the first principal components of each module in the network, in addition to the ROSMAP cohort specific trait of slope of cognitive decline, a Z score-scaled measure indicating degree of cognitive resilience of an individual compared to the mean age-dependent cognitive decline of the full ROSMAP cohort population^34, 35^. Correlation was performed using the WGCNA R package (v1.72-1) function plotEigengeneNetworks. For resilience PWAS enrichment of significance among PV-IN or CAMK2A neuron-enriched protein gene products (**Fig. 2F**), and for PV-IN 5xFAD DEPs (**Fig. 5O**), permutation-based enrichment of pooled significance from the PWAS was computed as previously published ()^36^, Software for this is available from https://www.github.com/edammer/MAGMA.SPA.

### Other sources of data used for analyses in this manuscript

MicroRNA affinity purification (miRAP) data from studies of PV-IN and Camk2a neurons was downloaded from supplemental information associated with the original miRAP publication^30^ and miRNA species with PV-IN vs. Camk2a neuronal enrichment patterns, were cross-referenced with predicted miRNA regulators in our PV-CIBOP and Camk2a-CIBOP studies.

### Other statistical considerations

Specific statistical tests used for individual experiments are detailed in the figure legends. Generally, all continuous variables were analyzed using parametric tests (two-tailed unpaired T test assuming equal variances when comparing 2 groups, or one-way ANOVA and post-hoc Tukey HSD tests for >2 group comparisons). Power calculations were not performed for individual experiments.

## Data availability

The mass spectrometry proteomics data generated by PV-CIBOP studies will be deposited to the ProteomeXchange Consortium via the PRIDE partner repository. Camk2a-CIBOP data can be obtained using dataset identifiers PXD027488 and PXD032161. The 2020 mouse Uniprot database (downloaded from https://www.uniprot.org/help/reference_proteome).

## Contributions

Conceptualization: PK, AMG, MJR, SR

Methodology and investigation: PL, AMG, CEG, BRT, JVS, AT, RSN, EBD, SM, LC, HX, DD, LBW, NTS, MJR, SR

Writing-Original draft: PK, AMG, MJR, SR

Writing-Review and Editing: PK, AMG, CEG, LBW, NTS, MJR, SR Funding acquisition: AMG, MJR, SR

Resources: NTS, MJR, SR

## SUPPLEMENTAL INFORMATION

### Supplemental Figures and legends

**Figure S1.**
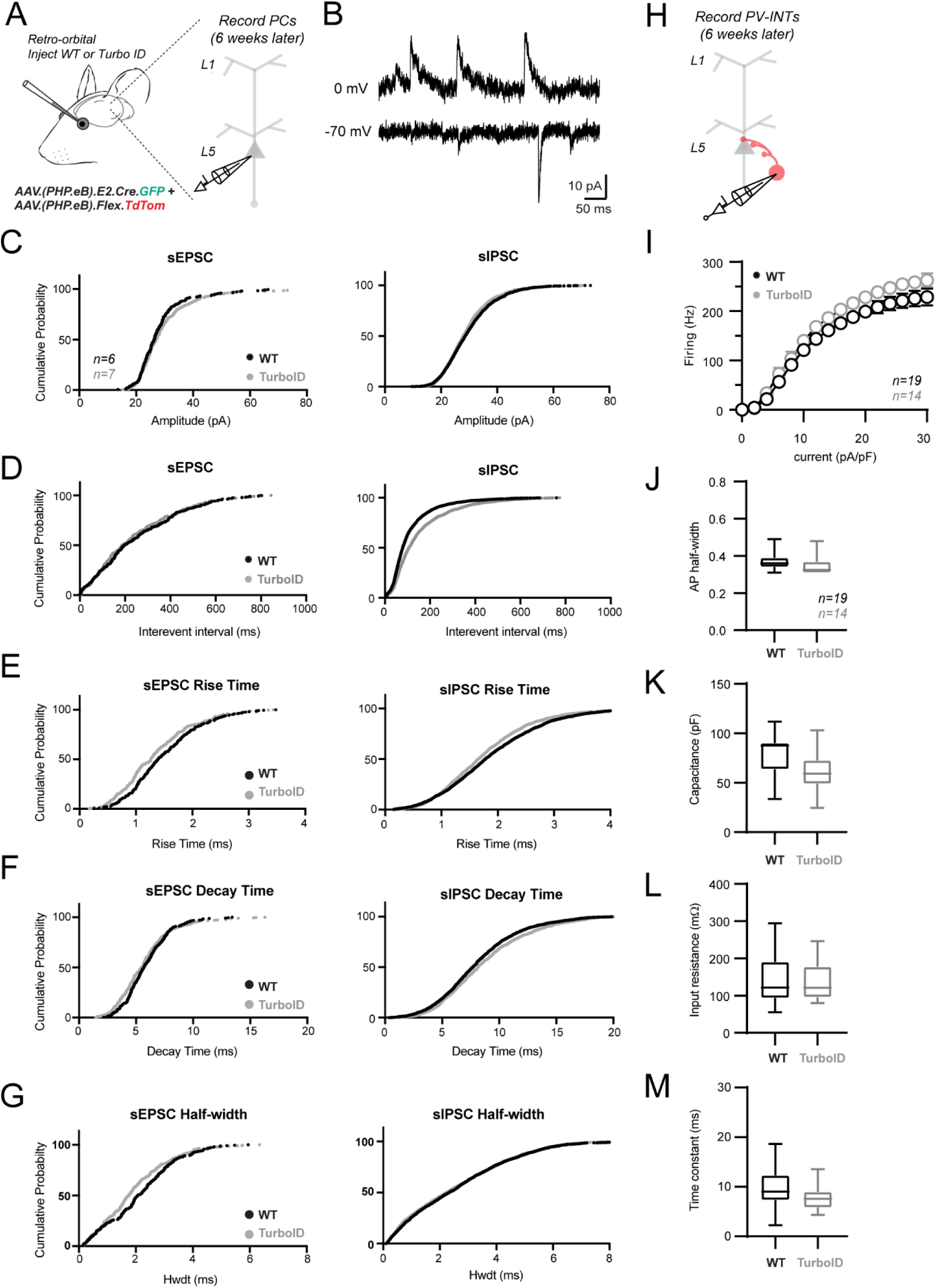
PV-CIBOP does not disrupt PV-IN or local circuit properties (related to Fig 1). **A.** During procurement of CIBOP tissues from RO-injected Rosa26^TurboID/wt^ (TurboID) and WT mice, a subset of the brain (SBFI region of S1 cortex) was used to immediately prepare acute slices for patch clamp recordings of unlabeled pyramidal neurons in layer 5. **B.** Example traces from a voltage clamp recording in a pyramidal neuron in layer 5 cortex. -70 and 0 mV holding potentials were interleaved throughout the recording to sample spontaneous EPSCs and IPSCs, respectively. **C-G.** Cumulative probability distribution curves for the amplitudes, frequency, and kinetic properties from all spontaneous EPSC and IPSC events recorded in pyramidal neurons of TurboID and WT mice. **H, I.** In the same experiments as depicted in **A** and **B**, fluorescent-targeted current clamp recordings were performed in TdTomato+ neurons as identified using combined video epifluorescent illumination. The GFP signal from the E2.Cre.GRP construct was not used, as it was generally much dimmer. Current injections (300 ms) of varying amplitude (0-30 pA/pF) were normalized to the individual cellular capacitance to control for potential variability between passive features. **J.** Narrow action potential widths at half-maximal amplitude (half-width) quantified in TdTomato+ between WT and Turbo ID mice. Half-width was measured from the 1^st^ spike elicited by current injection. Action potentials were generally ∼0.35 ms, characteristic of fast-spiking cortical PV-INs. **K-M.** Passive features measured in recordings from TdTomato+ PV-INs. For synaptic properties recorded in pyramidal neurons in **C-G,** average values from all spontaneous events from individual recordings were used for statistical analysis comparing TurboID (n=7) and WT (n=6) mice. All comparisons were p>0.05, unpaired two-tailed T-test. For PV-IN recording data in **J-M**, all TurboID (n=19) and WT (n=14) comparisons were p>0.05, unpaired two-tailed T-test.

**Figure S2.**
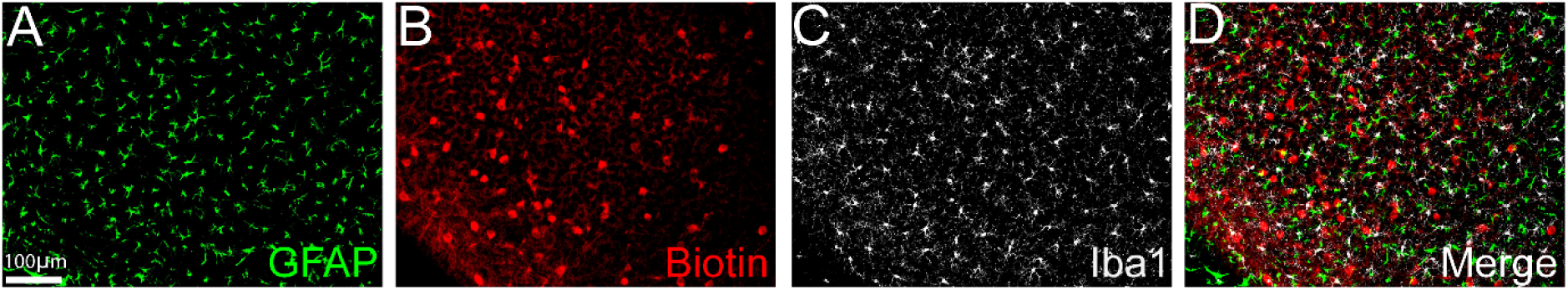
PV-CIBOP specifically labels PV-INs without off-target labeling of astrocytes or microglia (related to Fig 1). Representative immunofluorescence images (20x) from PV-CIBOP brain, confirming that biotinylation was not detected in astrocytes (GFAP+) microglia (Iba1+ positive) or astrocytes (GFAP+). Furthermore, microglial and astrocyte morphology was qualitatively similar in labeled and non-labeled animals.

**Figure S3.**
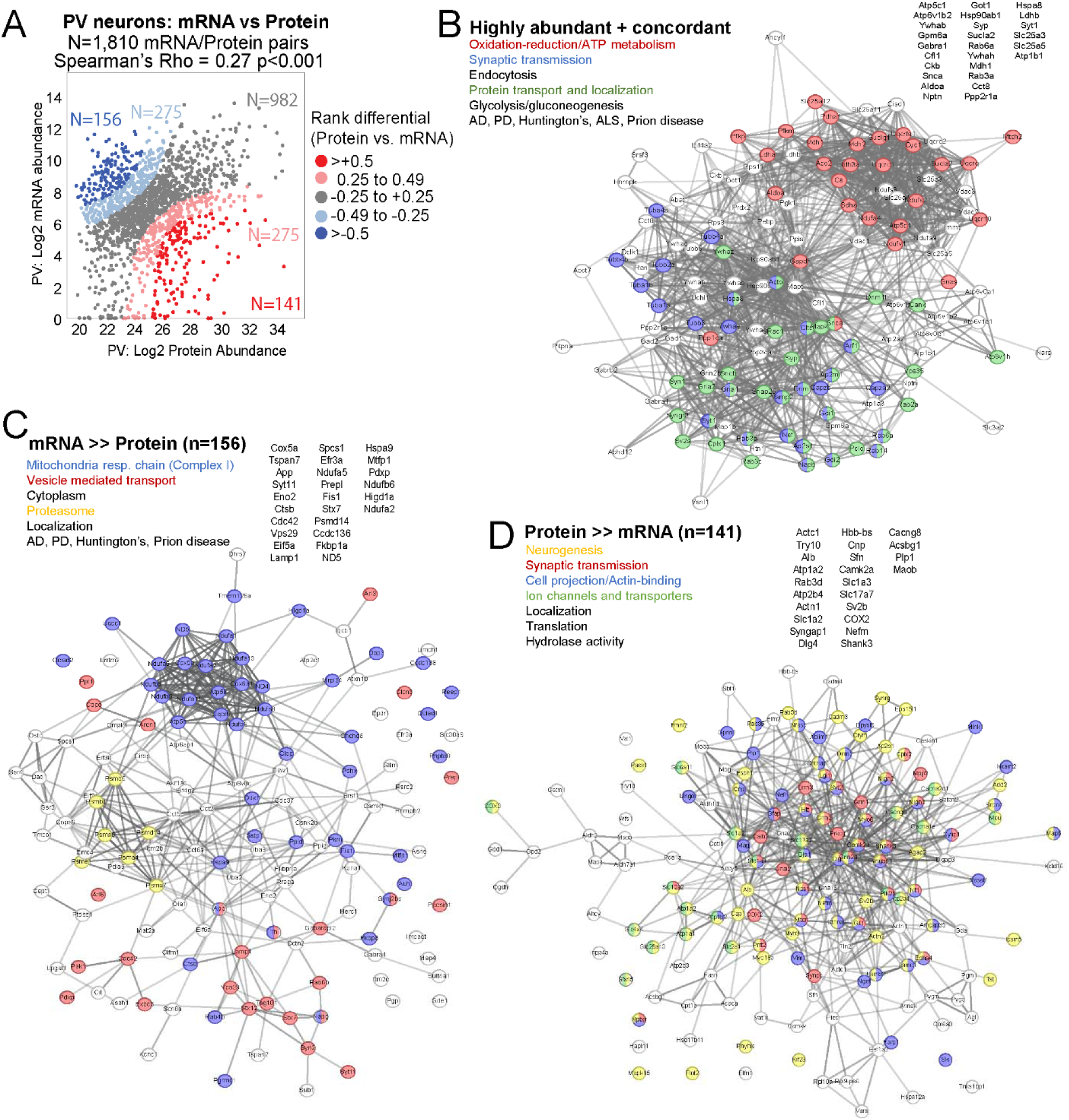
Assessment of protein-mRNA discordance in PV-INs (related to Fig 1M). **A.** Analysis of protein vs mRNA concordance in PV-INs, using PV-enriched proteins identified by PV-CIBOP and existing single nuclear transcriptomic data from the entire class of adult mouse PV-INs (Allen Brain Atlas). Based on differentials in rank abundances (protein vs. mRNA), discordant and concordant protein/mRNA pairs are highlighted. Spearman’s Rho for protein vs. mRNA correlation is shown. **B-D.** Three groups of protein/mRNA pairs were further analyzed: **B.** Group 1-Highly abundant and concordant: Protein rank differential between -0.25 and 0.49 and overall abundance >80^th^ percentile in both protein and mRNA datasets). **C.** Group 2: Discordance with mRNA greater than protein abundance (Protein vs. mRNA rank differential < -0.5). **D.** Group 3: Discordance with protein greater than mRNA abundance (Protein vs. mRNA rank differential > +0.5). Top gene symbols/protein IDs, top GSEA terms representative of each group, and STRING PPI diagrams (with colors indicating each ontology), are shown in each panel.

**Figure S4.**
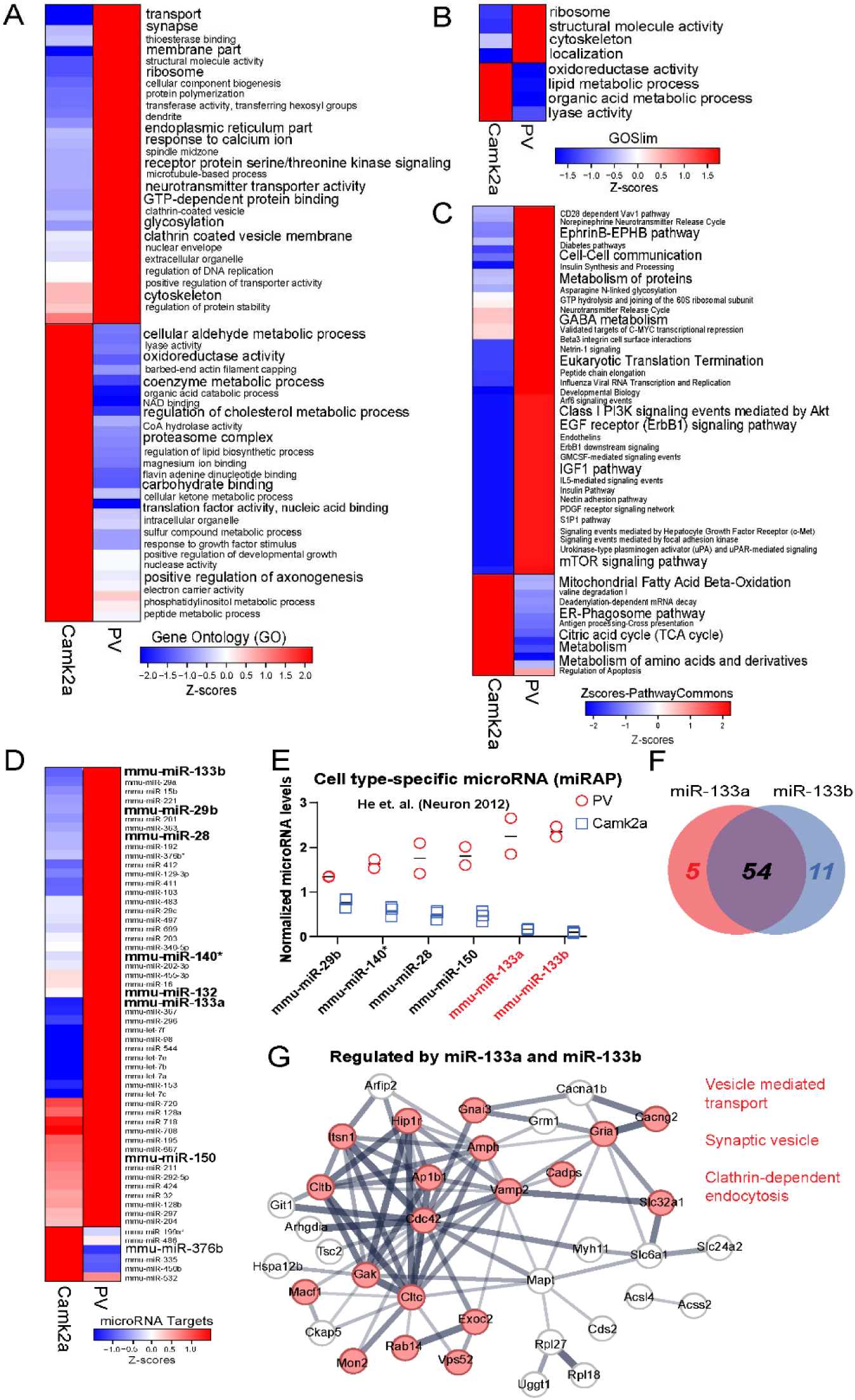
GSEA of PV-IN and Camk2a neuronal CIBOP proteomes (related to Fig 3). **A-D.** Heatmap representation of (**A**) Gene Ontology (GO) terms, (**B**) GOSlim terms, (**C**) PathwayCommons terms as well as (**D**) predicted upstream microRNA (miRNA) regulators that are over-represented in proteomic signatures of PV-INs and Camk2a neurons, using the CIBOP approach. **E**. Normalized miRNA levels as measured using the miRAP method (He et. al. Neuron 2012) that identified miRNAs that were differentially abundant in PV-INs and Camk2a neurons. The miRNAs enriched in PV-INs that are represented in this graph (particularly miR-133a and miR 133b), were also predicted as up-stream regulators of PV-IN proteomic signatures. Therefore, miR-133a and miR-133b are highly likely miRNA regulators that may be functionally important regulators of PV-INs. **F.** Venn Diagram representing shared and distinct target genes that are regulated by miR-133a and miR133-b in PV-IN proteomes. **G.** 54 proteins were identified as shared down-stream targets of both miR-133a and miR-133b in PV-INs, and these are represented as a STRING PPI network. Top GO terms emerging from this analysis include synaptic vesicle, vesicle transport and clathrin-dependent endocytosis terms. This indicates that miR-133a and miR-133b may be regulators of synaptic vesicle related function in PV-INs.

**Figure S5.**
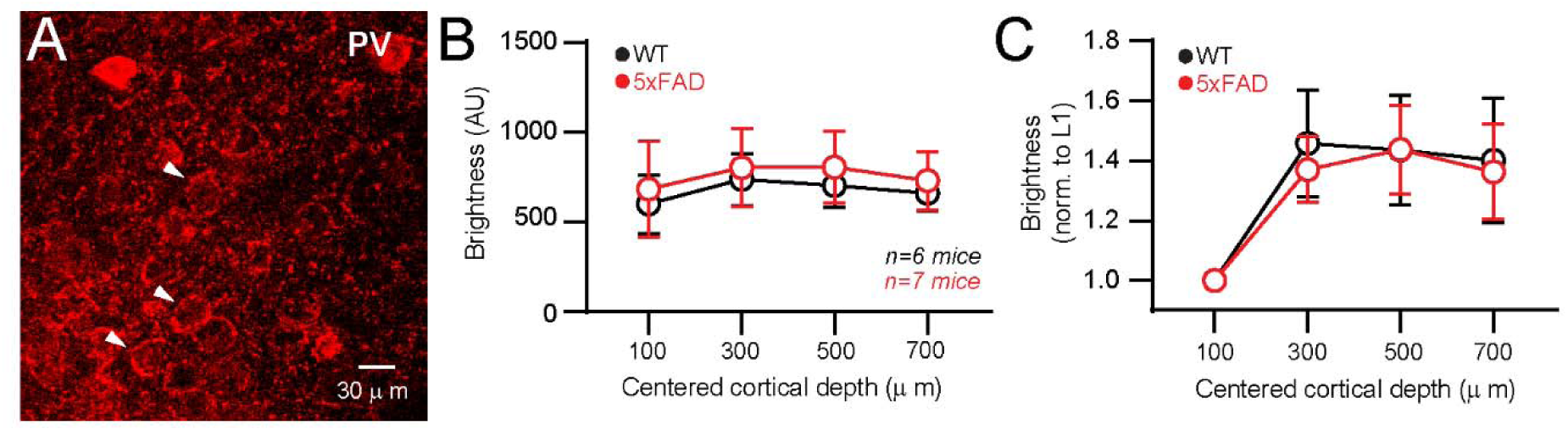
Parvalbumin IHC in WT and 5xFAD mice (related to Fig 4D-G). **A.** Representative IHC image of anti-parvalbumin staining in the L5 region of S1 cortex. White arrows denote examples of basket structures surrounding putative pyramidal and other neuron types. Extensive staining of PV-IN presynaptic basket structures was evident using this antibody staining method. **B.** Quantification of integrated fluorescence (in arbitrary units) from Alexa-594 secondary directed against Parvalbumin. Z-stacks were obtained from thin slices cortical WT and 5xFAD tissues. An ∼200 μm FOV 60X objective allowed for images centered at 100, 300, 500, and 700 (± 100) micron cortical depths. Background fluorescence measured at an offset location was subtracted from all images. **C.** Same quantification as in (B) but normalized to the FOV centered at 100 μm cortical depth. Reduced expression was apparent at this superficial depth, likely due to lack of extensive Parvalbumin-labeled structures with respect to deeper layers. Staining was consistent across all deeper areas corresponding to Layers 2/3, 4, and 5 and did not differ between WT and 5xFAD cortices.

**Figure S6.**
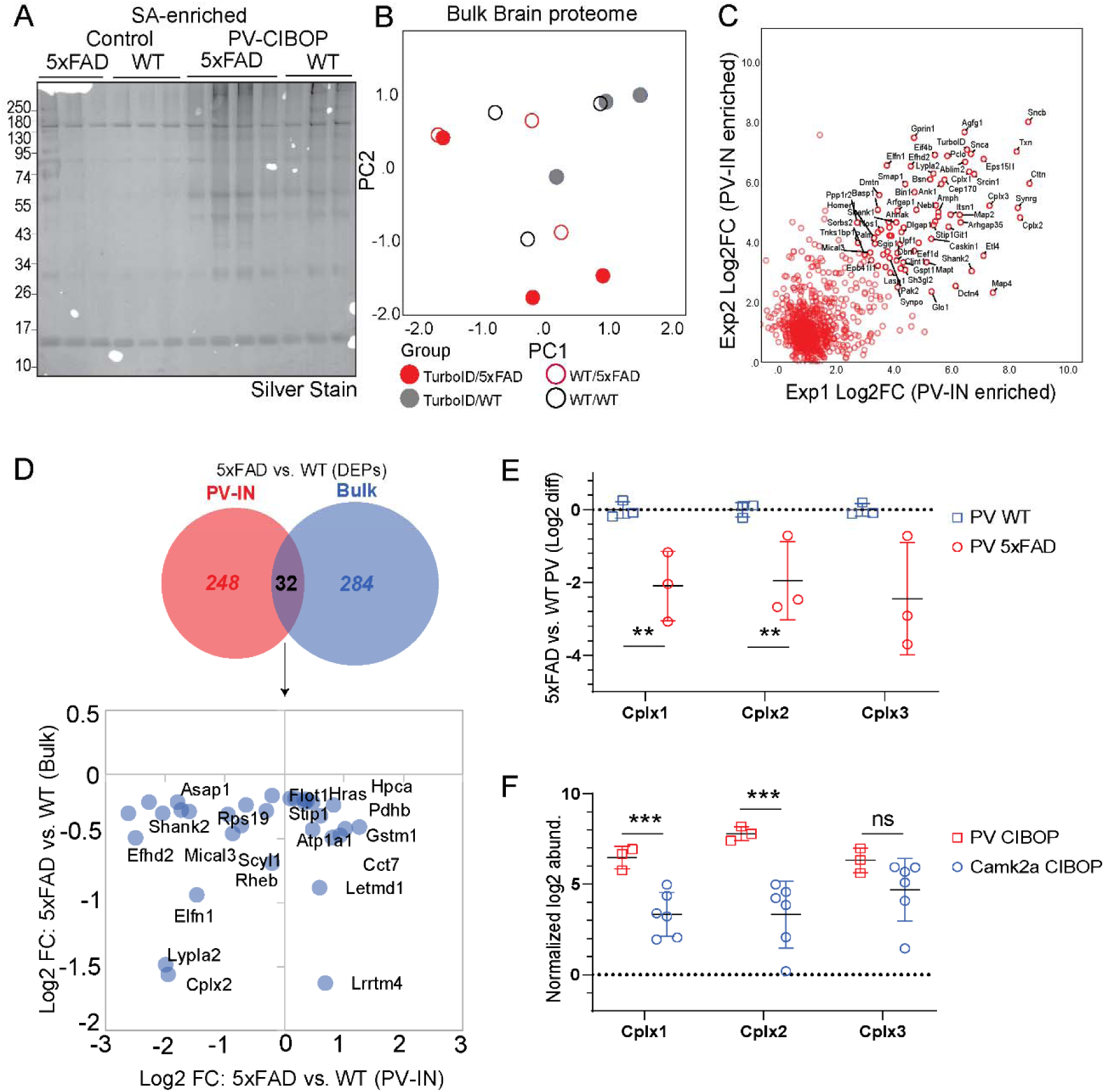
PV-CIBOP identifies novel proteomic changes occurring in PV-INs in early stages of A**β** pathology (related to Fig 5). **A.** Silver-stained gel of SA-enriched PV-IN proteins from experimental animals (in Fig 5) confirming higher protein enrichment from PV-CIBOP animals as compared to control animals. **B.** PCA of bulk brain (input) proteomes showing lack of an observable group-based clustering based on genotype (WT vs 5xFAD) or biotinylation (CIBOP vs controls). This contrasts with genotype differences observed in PV-IN proteomes presented in Fig 5. **C.** Level of agreement between two independent PV-IN proteomes using the CIBOP approach. Experiment 1: PV-CIBOP in WT mice presented in Fig 1. Experiment 2: PV-CIBOP in WT mice presented in Fig 5. Log2 fold changes (CIBOP vs control) of proteins that were labeled in both datasets, are shown. Overall correlation between two studies was moderate (Pearson’s Rho =0.61, p<0.001). Top PV-IN proteins identified by both studies, were similar (including Snca, Sncb, Cplx1, Cplx2, Cplx3, Elfn1, Bsn, as well as TurboID). **D.** DEPs (comparing 5xFAD vs. WT) identified at the level of the bulk proteome and PV-IN proteome were distinct except for 32 proteins (intersect). Agreement in level of differential abundances (log2 FC 5xFAD vs WT) between bulk and PV-IN proteomes was modest with the exception of some proteins (eg. Cplx2, Lypla2, Elfn1) which shown concordant decreased levels in both bulk and PV-IN proteomes. **E.** Protein levels of three complexins in the PV-IN proteome, comparing 5xFAD vs. WT genotypes (*p<0.05, **p<0.01, ***p<0.005). **F.** Protein levels of three complexins in PV-IN vs. Camk2a CIBOP proteomes (*p<0.05, **p<0.01, ***p<0.005).

**Figure S7.**
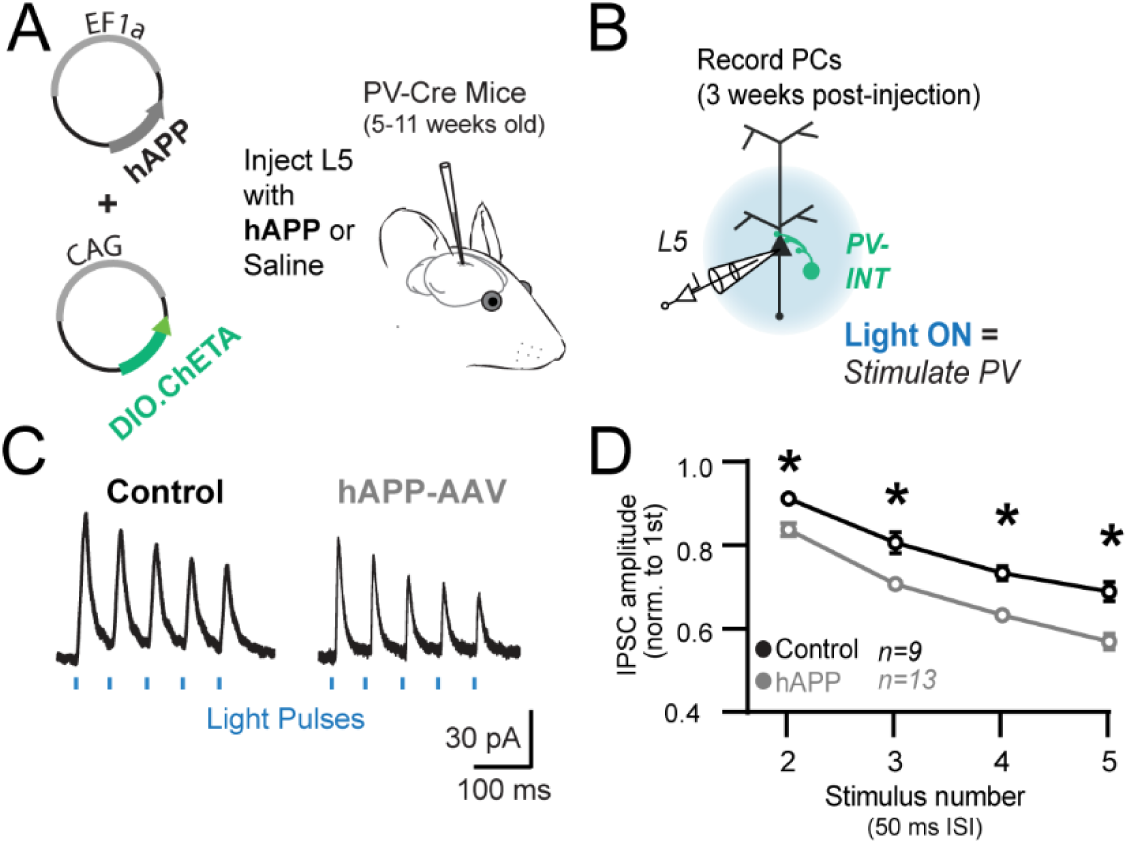
hAPP-AAV effect on PV neurotransmission (related to Figure 6). **A.** Experimental outline: PV-Cre were injected with AAV.CAG.DIO.ChETA and either AAV.EF1a.hAPP or saline control in the somatosensory cortex at 5-11 weeks of age. **B.** Three weeks post-injection, PV interneurons were stimulated at 20 Hz and nearby pyramidal cells were patched to examine the PV-PC paired-pulse ratio (PPR) and the multiple-pulse ratio (MPR). **C.** Example traces of optogenetically-evoked PV inhibitory post-synaptic currents on pyramidal cells for saline injected (left) and hAPP-AAV injected (right) cortices. **D.** IPSCs in AAV-hAPP injected mice displayed a significant change in synaptic dynamics as measured using MPR across all measured stimuli at 20 Hz. (*p<0.05 Two-way ANOVA with Sidak’s posthoc comparisons for each stimulus between hAPP and saline control experiments).

**Figure S8.**
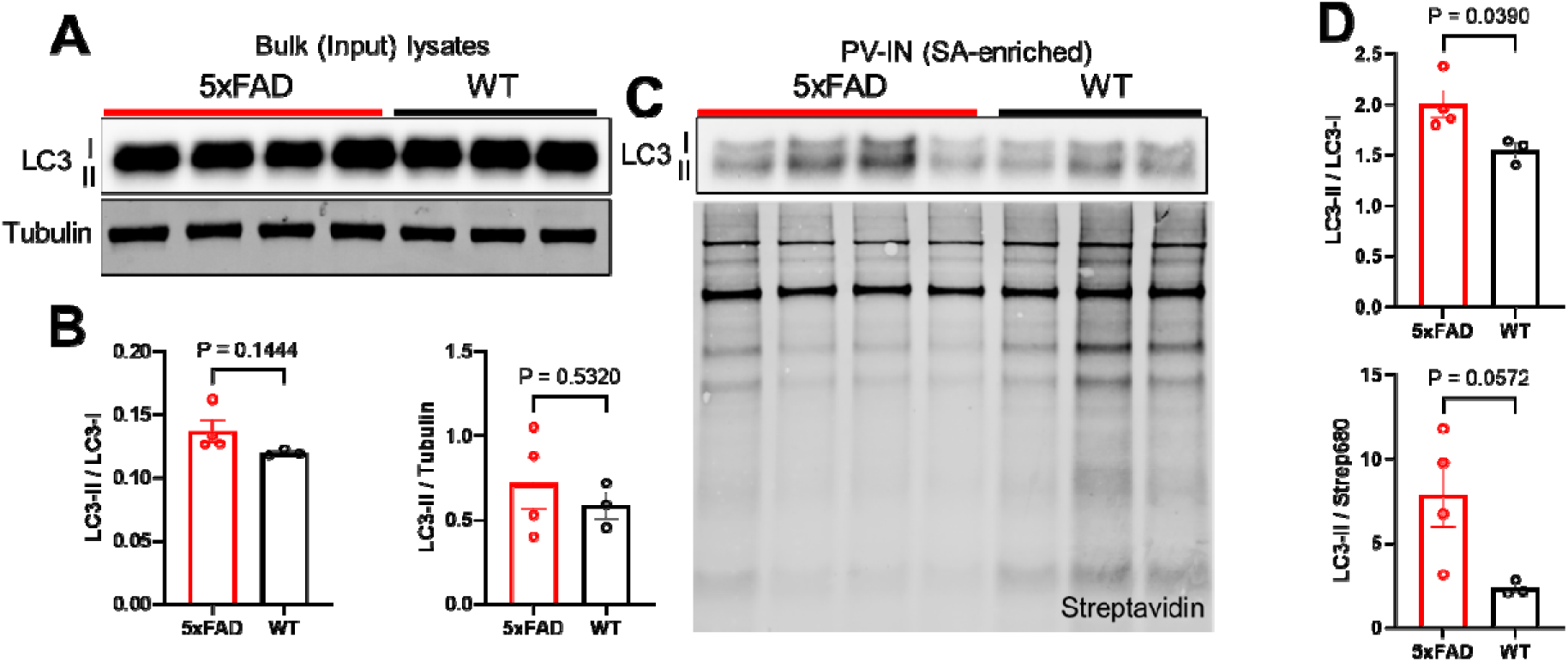
Effects of 5xFAD pathology on autophagic flux in PV-INs and bulk brain (related to Figure 7). **A.** Western blots of Input (bulk) brain lysates from WT/PV-CIBOP and 5xFAD/PV-CIBOP animals. Housekeeping protein used was tubulin. LC3-II and LC3-I were both detected, and LC3-II was normalized to either LC3-I (shown in Figure 7) or to tubulin. **B.** Quantification of LC3-II (normalized to LC3-I, or to alpha-tubulin) comparing 5xFAD to WT brain bulk lysates. P-values for two-sided unpaired T tests are indicated. **C.** Western blots of SA-enriched biotinylated proteins (PV-IN proteins) from WT/PV-CIBOP and 5xFAD/PV-CIBOP animals. LC3-II and LC3-I were both detected, and LC3-II was normalized to either LC3-I (shown in Figure 7) or to total biotinylation signal (streptavidin). **D.** Quantification of LC3-II (normalized to LC3-I [top] or to streptavidin [bottom]) comparing 5xFAD to WT brain. P-values for two-sided unpaired T tests are indicated.

**Supplemental Datasheets:**

**Datasheet 1.** LFQ-MS data related to Fig 1: PV-IN proteome

**Datasheet 2.** LFQ-MS data related to Fig 2: PV-IN vs Camk2a CIBOP proteome

**Datasheet 3.** Neuronal subtype marker lists used for analyses in Fig 3

**Datasheet 4.** Mouse TMT-MS data related to Fig 4.

**Datasheet 5**. PV-CIBOP in 3 mo 5xFAD and WT mice: Bulk and SA-enriched proteomes and associated analyses, related to Fig 5

**Datasheet 6.** Analysis of protein-half life distribution in PV-IN DEPs (5xFAD vs. WT CIBOP), related to Fig 7

**Datasheet 7.** List of reagents including antibodies

